# Estimating gene expression and codon specific translational efficiencies, mutation biases, and selection coefficients from genomic data alone

**DOI:** 10.1101/009670

**Authors:** Michael A. Gilchrist, Wei-Chen Chen, Premal Shah, Cedric L. Landerer, Russell Zaretzki

## Abstract

Extracting biologically meaningful information from the continuing flood of genomic data is a major challenge in the life sciences. Codon usage bias (CUB) is a general feature of most genomes and is thought to reflect the effects of both natural selection for efficient translation and mutation bias. Here we present a mechanistically interpretable, Bayesian model (ROC SEMPPR) to extract biologically meaningful information from patterns of CUB within a genome. ROC SEMPPR, is grounded in population genetics and allows us to separate the contributions of mutational biases and natural selection against translational inefficiency on a gene by gene and codon by codon basis. Until now, the primary disadvantage of similar approaches was the need for genome scale measurements of gene expression. Here we demonstrate that it is possible to both extract accurate estimates of codon specific mutation biases and translational efficiencies while simultaneously generating accurate estimates of gene expression, rather than requiring such information. We demonstrate the utility of ROC SEMPPR using the *S. cerevisiae* S288c genome. When we compare our model fits with previous approaches we observe an exceptionally high agreement between estimates of both codon specific parameters and gene expression levels (*ρ >* 0.99 in all cases). We also observe strong agreement between our parameter estimates and those derived from alternative datasets. For example, our estimates of mutation bias and those from mutational accumulation experiments are highly correlated (*ρ* = 0.95). Our estimates of codon specific translational inefficiencies are tRNA copy number based estimates of ribosome pausing time (*ρ* = 0.64), and mRNA and ribosome profiling footprint based estimates of gene expression (*ρ* = 0.53 *−* 0.74) are also highly correlated, thus supporting the hypothesis that selection against translational inefficiency is an important force driving the evolution of CUB. Surprisingly, we find that for particular amino acids, codon usage in highly expressed genes can still be largely driven by mutation bias and that failing to take mutation bias into account can lead to the misidentification of an amino acid’s ‘optimal’ codon. In conclusion, our method demonstrates that an enormous amount of biologically important information is encoded within genome scale patterns of codon usage, accessing this information does not require gene expression measurements, but instead carefully formulated biologically interpretable models.

## Introduction

Genomic sequences encode a trove of biologically important information. Over 49,600 genomes are currently available from the Genomes OnLine Database (Pagani *et al*., 2012) alone and the flow of newly sequenced genomes is expected to continue far into the future. As a result, developing ways to turn this data into useful information is one of the major challenges in the life sciences today. Although great strides have been made in extracting this information, ranging from the simple, e.g. identification of protein coding regions, to the more difficult, e.g. identification of regulatory elements (Hughes *et al*., 2000; Wasserman and Sandelin, 2004; Dunham *et al*., 2012; Kundaje *et al*., 2015), much of this information remains untapped. To address one aspect of this challenge, we present a method to estimate the expression levels of every gene, codon specific selection coefficients, and mutation biases *solely* from patterns of codon usage bias (CUB) in protein coding sequences within a genome.

One of the earliest arguments against neutrality between synonymous codon usage was given by Clarke (1970). Since then, evidence for selection acting on CUB has been repeatedly observed. CUB clearly varies systematically within and between open reading frames (ORFs) within a species as well as across species (Grantham *et al*., 1980; Ikemura, 1981a, 1985; Bennetzen and Hall, 1982; Sharp and Li, 1987; Andersson and Kurland, 1990b; Qin *et al*., 2004; Gilchrist and Wagner, 2006; Chamary *et al*., 2006; Hershberg and Petrov, 2008; Plotkin and Kudla, 2011). These patterns in CUB are driven by two evolutionary forces: mutation bias and natural selection (Ikemura, 1981a; Bulmer, 1988, 1991). Current evidence supports multiple selective forces contributing to the evolution of CUB. Most of these hypothesized selective forces affect the efficiency and efficacy of ORF translational through factors such as ribosome pausing times (Andersson and Kurland, 1990b; Bulmer, 1991; Sørensen and Pedersen, 1991; Plotkin and Kudla, 2011; Shah and Gilchrist, 2011a), missense and nonsense errors (Kurland, 1987, 1992; Akashi, 1994; Gilchrist, 2007; Drummond and Wilke, 2008, 2009), co-translational protein folding (Thanaraj and Argos, 1996; Kimchi-Sarfaty *et al*., 2007; Tsai *et al*., 2008; Pechmann and Frydman, 2013), equalizing tRNA availability (Qian *et al*., 2012), and the stability and/or accessibility of mRNA secondary structures (Kudla *et al*., 2009; Tuller *et al*., 2010; Gu *et al*., 2012; Bentele *et al*., 2013). The relative importance of each of these selective forces is expected to vary both within and between genes. The effects of these forces can be unified within a single framework by considering how the codon usage of a given ORF alters the ratio of the expected cost of protein synthesis over the expected benefit of protein synthesis, or the cost-benefit ratio *η* for short (Gilchrist *et al*., 2009) (see Methods).

One likely way different synonymous codons lead to changes in a gene’s cost-benefit ratio *η* results from differences in the abundances of cognate and near cognate tRNAs and the stability of the Watson-Crick base pairing between a given codon and tRNA anticodons (Ikemura, 1981a; Zaher and Green, 2009; Plotkin and Kudla, 2011). These differences, in turn, are predicted to lead to differences in ribosome pausing times and error rates between codons. Specifically, codons with higher abundances of cognate and near-cognate tRNAs are thought to have both shorter pausing times and lower error rates than codons with lower abundances of cognate and near-cognate tRNA (Ikemura, 1981a; Kurland, 1992, though see Shah and Gilchrist (2010) for a more nuanced view).

The assumption that natural selection favors codon usage which reduces the protein synthesis cost-benefit ratio *η* implies that the strength of this selection should scale with the gene’s protein synthesis rate: highly expressed genes should show the strongest bias for codons with shorter pausing times and error rates (Ikemura, 1981a, 1985; Sharp and Li, 1986, 1987). As a result, given sufficiently large *N*_*e*_, such that high expression genes contain some signal of adaptation, the patterns of CUB observed within a genome should contain a significant amount of information about the average protein synthesis rate *ϕ* for a given gene. Further, because low expression genes are under very weak selection to reduce *η*, their patterns of CUB should provide information on the mutational biases experienced within a genome.

Accessing this information held within CUB patterns of an organism’s genome has been the focus of several decades of research in molecular evolution. However, most approaches examine mutation bias and selection in isolation and, ignore their possible interactions. The strength of mutation bias has typically been investigated by comparing the differences in GC content of synonymous sites of codons to the rest of the gene (Galtier *et al*., 2001; Knight *et al*., 2001; Palidwor *et al*., 2010). Numerous methods have been used to quantify or describe selection on synonymous codon usage.

For example, Sharp and Li (1987) relied on the codon usage in a set of highly expressed genes to identify the ‘optimal’ codon for a given amino acid as these genes are under stronger selection to be translated efficiently and accurately. Approaches that focus on a subset of high expression genes in this way implicitly assume the contribution of mutation bias to CUB is overwhelmed by natural selection and, therefore, can be ignored. As our results show, because this view lacks an explicit population genetics framework it is likely overly simplistic and may lead to the misidentification of ‘optimal’ codons.

Phylogeny based models of protein evolution, some of which are derived from population genetics models, have also been used to generate estimates of codon-specific selection coefficients and mutation biases (Tamuri *et al*., 2012; Rodrigue *et al*., 2010; Yang and Nielsen, 2008). Other approaches have relied on intra-specific variation to make similar types of inferences (Keightley and Eyre-Walker, 2007; Lawrie *et al*., 2013) or a combination of inter-specific divergence and intra-specific variation Akashi (1995). However, all of these approaches fail to disentangle how the contributions of mutation bias and natural selection change with gene expression. Furthermore, these models are either fitted independently across genes and thus estimate a large number of gene specific parameters from a relatively small amount of data or assume that the magnitude of selection is uniform across genes.

We, along with others, have previously worked to link gene expression levels to patterns of CUB by nesting a mechanistic model of protein translation into a population genetics model of allele fixation in order to estimate codon specific mutation and selection parameters (Gilchrist and Wagner, 2006; Gilchrist, 2007; Shah and Gilchrist, 2011a; Wallace *et al*., 2013). Although these methods represent significant advances in estimating codon specific mutation biases and selection coefficients from genomic data, they are limited to genomes with independent measurements of gene specific protein synthesis rates or a close proxy. Historically, mRNA abundances have been used as such a proxy due to the fact that generating reliable genome scale measurements of protein synthesis is an expensive undertaking (Arava *et al*., 2005; Ingolia *et al*., 2009a; Li *et al*., 2014, e.g). In contrast, the method proposed here does away with the necessity of having protein synthesis rate estimates (or their proxy) and *provides* estimates of the average protein synthesis rate for each gene, *ϕ*. Importantly, our method also provides estimates of codon specific mutation biases and translational inefficiencies, which is the additive contribution of a codon to the cost of protein synthesis.

Furthermore, we can combine our gene specific estimates of protein synthesis rates and codon specific translational inefficiencies to produce estimates of the strength of natural selection on synonymous substitutions on a gene by gene and codon by codon basis. Estimating gene-specific selection coefficients on synonymous codons is critical to determining whether a gene is evolving under purifying or positive selection. Current models to identify the selection regime under which a gene evolves rely on estimates of the rates of non-synonymous changes to rates of synonymous changes (*dN/dS*) (Li *et al*., 1985; Nei and Gojobori, 1986; Yang and Nielsen, 2000). However, the commonly made assumption that all synonymous changes within a gene are neutral can bias values of *dN/dS* towards over-estimating the number of genes evolving under positive selection (Spielman and Wilke, 2015). By accurately estimating strength of selection on synonymous changes, researchers can begin to explicitly incorporate these effects into methods for identifying purifying and positive selection.

In order to extract information from the genome wide patterns of CUB using our Stochastic Evolutionary Model of Protein Production Rate (SEMPPR) (Gilchrist, 2007; Shah and Gilchrist, 2011a), we build on the Bayesian statistical advances of Wallace *et al*. (2013). Because the costs in our model can be interpreted as proportional differences in ribosome overhead costs (ROC) due to ribosome pausing, for simplicity we refer to the model formulated here as ROC SEMPPR (see Methods).

Using the *Saccharomyces cerevisiae* S288c genome as an example, we demonstrate that ROC SEMPPR can be used to accurately estimate differences in codon specific mutation biases and contributions to the protein synthesis cost-benefit function *η without* the need for gene expression data. ROC SEMPPR’s codon specific estimates of mutation biases and translational inefficiencies generated without gene expression data match almost exactly those generated with gene expression data (Pearson correlation coefficient *ρ >* 0.99 for both sets of parameters). In the end, we observe a Pearson correlation coefficient of *ρ* = 0.72 between our predicted protein synthesis rates and the mRNA abundances from Yassour *et al*. (2009) (which was identified as the most reliable dataset out of five different mRNA abundance datasets by Wallace *et al*. (2013)).The variation between our predictions and Yassour *et al*. (2009)’s measurements is on par with the variation observed between mRNA abundance measurements from different laboratories (Wallace *et al*., 2013). Further, our predictions show strong and significant correlations with measurements of mRNA abundance from four other labs and estimates of protein synthesis rates based on ribosome profiling data from three other labs (Supporting Figures S4 and S5).

By releasing our work as a stand alone package in R (see Chen *et al*. (2014)), researchers can potentially take the genome of any microorganism and obtain accurate, quantitative information on the effect of synonymous substitutions on protein translation costs, gene expression levels, and the strength of selection on codon usage bias.

## Results

The posterior means estimated from our Bayesian MCMC simulation of ROC SEMPPR demonstrate two key facts: 1) we are able to estimate the strength of selection on synonymous codon usage bias from the patterns of codon usage observed within a genome and, 2) we can attribute this selection to the interaction of two underlying biological traits: difference between synonymous codons in their contribution to the cost-benefit ratio *η* for protein synthesis and the protein synthesis rate of the ORF *ϕ* averaged across its various environments and lifestages.

For this study, we scale our codon specific translational inefficiencies relative to the strength of genetic drift, 1*/Ne,*

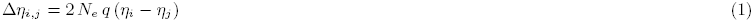

where *q* described the proportional decline in fitness per ATP wasted per unit time. More specifically, Δ*η_i,j_* describes the difference in the contribution of synonymous codons *i* and *j* to the protein synthesis cost benefit-ratios of an ORF, (*ηi − ηj*), scaled by effective population size *Ne* ≫ 1 and the relative fitness cost of expending an extra ATP per unit time, *q*. The greater the contribution of a codon to *η*, the greater its inefficiency. For a set of synonymous codons, by convention, we define codon 1 as the codon with the lowest inefficiency, i.e. the codon which makes the smallest additive contribution to *η* and is most favored by selection. Thus, Δ*η*_*i,*1_ = 0 for *i* = 1 and Δ*η*_*i,*1_ *>* 0 for *i >* 1. For notational simplicity, we will only include the subscripts when needed for clarity.

At equilibrium, under the weak-mutation regime (Sella and Hirsh, 2005b; Shah and Gilchrist, 2011b; McCandlish and Stoltzfus, 2014), the expected frequency of observing a synonymous codon *i* (*p*_*i*_) of an amino acid in a gene with an average protein synthesis rate *ϕ* follows a multinomial logistic distribution. Specifically, for a given amino acid *a* with *n*_*a*_ unique codons

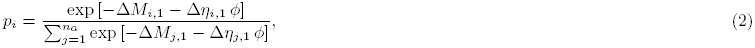

where Δ*M*_i,1_ is a unitless measure of codon specific mutation bias. Note that, as with Δ*η*, Δ*M*_*i,*1_ = 0 for *i* = 1 but, unlike with Δ*η*, Δ*M*_*i,*1_ can be positive or negative. Further, because it relies on the stationary probability of observing a synonymous codon, ROC SEMPPR can only detect variation in mutation bias, not variation in absolute mutation rates. Additional model details can be found in the Methods and Materials.

The utility of Equation (2) is that it allows us to probabilistically link ROC SEMPPR’s parameters of interest, i.e. codon specific differences in mutation biases, 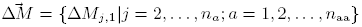, translational inefficiencies 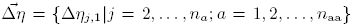, and gene specific protein synthesis rates, 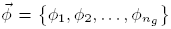, to the CUB patterns observed within and between ORF of a given genome. The terms *n*_aa_ represents the number of amino acids that use multiple codons while *n*_*g*_ represents the number of genes in the genome, respectively.

Because moving between the synonymous codon groups (TCA, TCC, TCG, TCT) and (AGC, AGT) for Ser requires at least one non-synonymous nucleotide substitution, we treated these two groups as if they were different amino acids, Ser_4_ and Ser_2_, respectively. So while strictly speaking, 18 of the canonical 20 amino acids use more than one codon, because we treat Ser as two separate amino acids, Ser_2_ and Ser_4_, for our purposes *n*_aa_ = 19. Assuming a log-normal distribution (LogN) with a mean of 1 as the prior for *ϕ* allows us to employ a random walk Metropolis chain to estimate the posteriors for 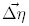, 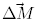, and 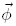 without the need for any laboratory measurements of gene expression, 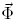. This ability to fit our ROC SEMPPR model *without* 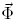 data is the major advance of our work over Wallace *et al*. (2013). Tables with estimates of gene specific protein synthesis rates 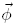, mutation biases, Δ*M*, and translational inefficiencies, Δ*η*, based on ROC SEMPPR’s posterior sampling for the *S. cerevisiae* genome can be found in the Supporting Materials.

### Evaluating Model Parameter Estimates

Briefly, when fitted to the *S. cerevisiae* S288c genome, we find nearly perfect agreement between ROC SEMPPR’s *with* and *without 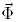* estimates for codon specific protein synthesis translational inefficiencies, Δ*η*, and mutation bias, Δ*M* (Pearson correlation *ρ* > .99 for both sets of parameters, see Figures 1) and 2). We note that, with the exception Arginine’s Δ*η*CGT,AGA, the central 95% Credibility Intervals (CIs) for ROC SEMPPR’s Δ*η* and Δ*M* parameters do not overlap with zero (see Supplemental Tables S1-S4). These results indicate that information on the genome scale parameters,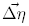 and 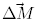 are robustly encoded and estimable from CUB patterns and that 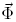, provides little additional information.

**Figure 1:**
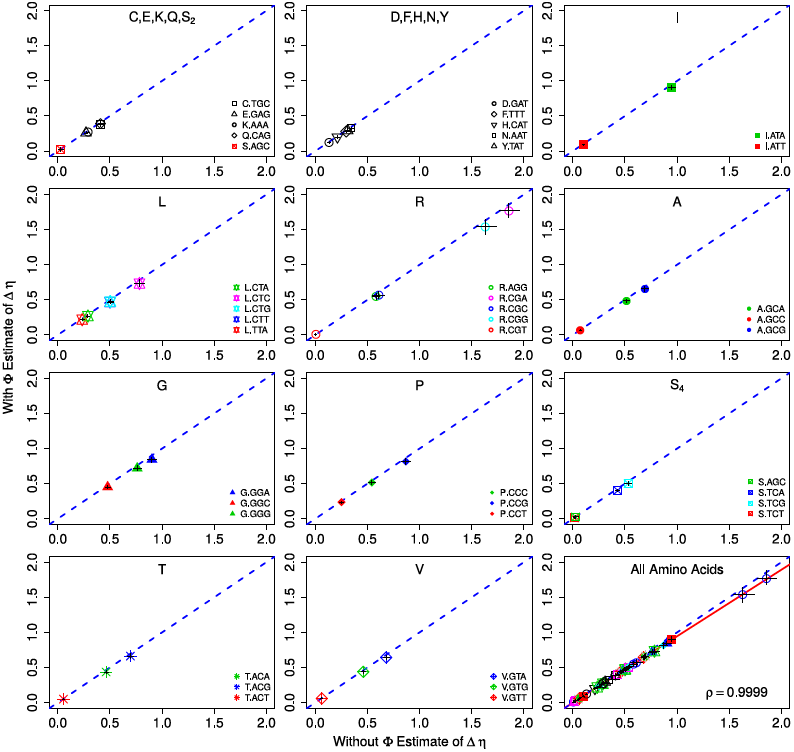
Comparison of *with* and *without 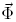* ROC SEMPPR estimates for codon specific differences in translational efficiencies Δ*η* which have units 1*/*(*t* protein) where the units of time are set such that the average protein synthesis rate across the genome, 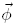 equals 1. To improve legibility of the plots the two codon amino acids have been combined into two plots and all of the amino acids with *>* 2 codons into separate plots. The dashed blue line represents the 1:1 line between axes and error bars indicate the 95% posterior credibility intervals (CIs) for each parameter. For both the *with* and *without 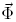* fits of ROC SEMPPR, all codons but one, Arg codon CGT, have CIs that do not overlap with 0. As illustrated in the last plot, a linear regression between estimates of Δ*η* for all codons produces a correlation coefficient *ρ >* 0.999.

**Figure 2:**
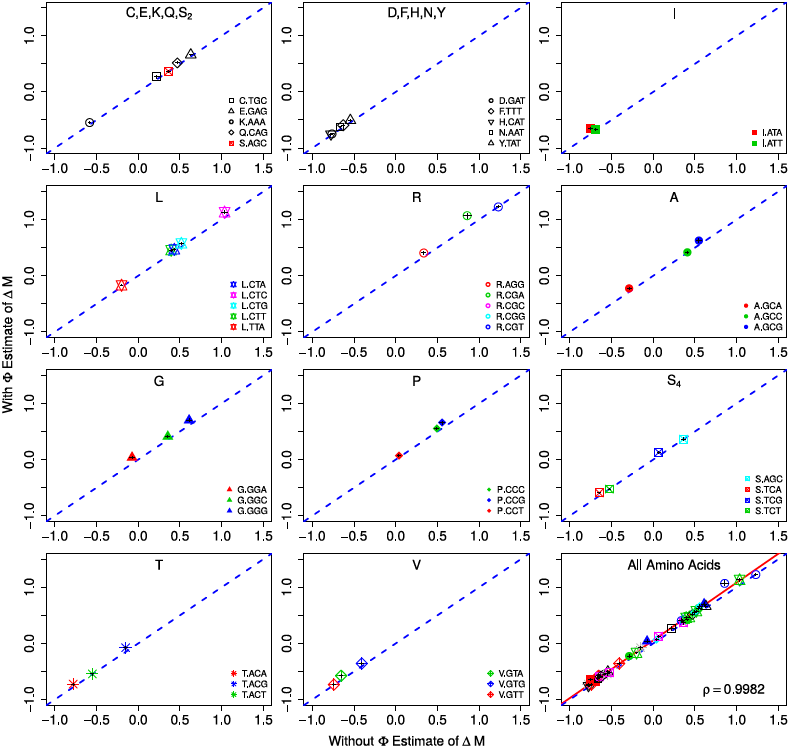
Comparison of *with* and *without 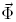* ROC SEMPPR estimates for codon specific differences in mutation biases terms Δ*M* which are unitless. Specifically, Δ*M*_*i,*1_ equals the natural logarithm of the ratio of the frequencies of synonymous codon 1 to *i* in the absence of natural selection. To improve legibility of the plots the two codon amino acids have been combined into two plots and all of the amino acids with *>* 2 codons into separate plots. The dashed blue line represents the 1:1 line between axes and error bars indicate the 95% posterior credibility intervals (CIs) for each parameter. For both the *with* and *without* fits of ROC SEMPPR, all codons have CIs that do not overlap with 0. As illustrated in the last plot, a linear regression between estimates of Δ*M* for all codons produces a correlation coefficient *ρ >* 0.998.

**(Approximate Location of Figure 1)**

**(Approximate Location of Figure 2)**

Instead of simply comparing our ROC SEMPPR model’s *without 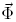* estimates of Δ*M* and Δ*η* to its *with 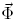* estimates, we can also compare these parameters to other data. Due to the detailed balance requirement of the stationary distribution of our population genetics model (Sella and Hirsh, 2005a), differences in Δ*M* values between codons that can directly mutate to one another will equal the log of the ratio of their mutation rates. Thus, our estimates of Δ*M* provide testable hypotheses about the ratio of mutation rates in *S. cerevisiae*. We use estimates of per base-pair mutation rates from a recent high-throughput mutation accumulation experiment in *S. cerevisiae* (Zhu *et al*., 2014). These experimental estimates of mutation bias, Δ*M ^e^*, are calculated as

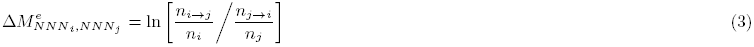

Where 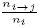is the number of *i → j* mutations observed per *n*_*i*_ bases in the genome. Since mutations in mutation accumulation experiments are strand agnostic, i.e. they do not distinguish between the coding and template strand nucleotides, we cannot distinguish between the mutations NNC*→*NNG and NNG*→*NNC nor NNA*→*NNT and NNT*→*NNA. As a result, our empirical estimates of 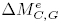 and 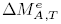 are set to 0. We find that our estimates of codon specific mutation rates correlate highly with empirical mutation rates in *S. cerevisiae* (*ρ* = 0.95, Figure 3).

**Figure 3:**
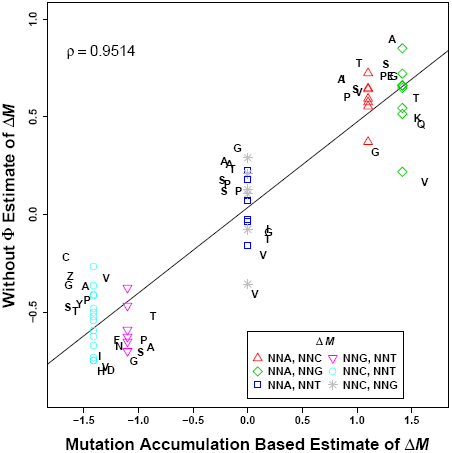
Comparison of *without 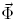* estimates of codon specific mutation biases Δ*M* and estimates generated from mutation accumulation experiments (Zhu *et al*., 2014). For each amino acid the codon with the shortest pausing time is used as a reference and are not shown because, by definition their Δ*M* values are 0. Pearson correlation coefficient *ρ* for all of the codons is given. The solid line represents the best fit linear regression.

**(Approximate Location of Figure 3)**

Unlike mutation bias parameters, empirical estimates of the codon specific differences in translational efficiencies do not exist. However, one of the simplest ways of linking a codon to *η* is based on the indirect cost of codon specific ribosome pausing during translation. That is, *η_i_ − η_j_ ∞ t_i_ − t_j_* where *t_i_* is the average time a ribosome pauses when translating codon *i*. We calculate empirical estimates of pausing times based on a simple model of translation where pausing times at a codon depend only on its cognate tRNA abundances and associated wobble parameters (Ikemura, 1981b; Andersson and Kurland, 1990a; Sørensen and Pedersen, 1991; Kanaya *et al*., 1999; Gilchrist and Wagner, 2006; Zaher and Green, 2009; Shah *et al*., 2013).

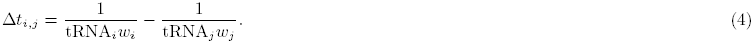

Specifically, tRNA_*i*_ is the gene copy number of the tRNA that recognize codon *i* and *w_i_* is the wobble penalty between the anti-codon of tRNA_*i*_ and codon *i*. When a codon is recognized by its canonical tRNA, we set *w*_*i*_ = 1. We assume a purine-purine (RR) or pyrimidine-pyrimidine (YY) wobble penalty to be 39% and a purine-pyrimidine (RY/YR) wobble penalty to be 36% based on Curran and Yarus (1989) and Lim and Curran (2001). We find that our genome-wide estimates of Δ*t* are positively correlated with empirical estimates of Δ*t* in *S. cerevisiae* (*ρ* = 0.64, Figure 4).

**Figure 4:**
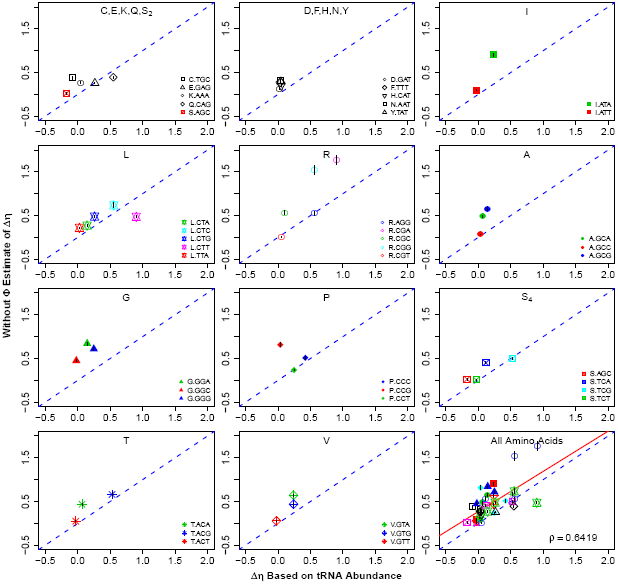
Comparison of *without 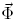* estimates of codon specific translational inefficiencies Δ*η* and estimates of differences in ribosome pausing times, Δ*t* based on tRNA gene copy number and wobble inefficiencies. For each amino acid the codon with the shortest pausing time is used as a reference and are not shown because, by definition their Δ*η* values are 0. Pearson correlation coefficient *ρ* for all of the codons is given. The dashed blue line represents the 1:1 line and the red line represents the best fit linear regression line.

**(Approximate Location of Figure 4)**

### Predicting Protein Synthesis Rates

Given the strong correlation between ROC SEMPPR’s *with* and *without 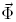* estimates of the codon specific mutation biases 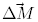 and translational inefficiencies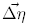, it is not surprising that *with* and *without 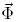* estimates of *ϕ* from ROC SEMPPR are highly correlated (*ρ* = 0.99, Figure 5(a)). More importantly, the *without 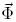* based estimates of *ϕ* show substantial correlation with the mRNA abundance based estimates of 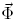 values from Yassour *et al*. (2009) (*ρ* = 0.72, Figure 5 (b)). To be clear, these 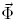 values are the same values used as inputs to the *with 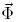* model fits.

Supporting Figures S4 and S5 explore this issue further by plotting ROC SEMPPR’s posterior mean estimates of *ϕ* produced *with* and *without 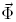* against eight sets of experimental data. This data includes three genome wide estimates based on ribosome-profiling (RPF) measurements (Ingolia *et al*., 2009b; Artieri and Fraser, 2014; McManus *et al*., 2014) and five other genome wide estimates of mRNA abundances (Arava *et al*., 2003; Nagalakshmi *et al*., 2008; Holstege *et al*., 1998; Sun *et al*., 2012). The *with 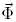* posterior estimates are generated using mRNA abundance measurements from Yassour *et al*. (2009) and are, therefore, independent of the measurements from other labs. Correlation between *ϕ* estimates for the *without 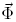* ROC SEMPPR fits and measured mRNA abundances range from 0.534 to 0.707, and measured RPF reads range from 0.629 to 0.742. The correlation between *ϕ* estimates for the *with 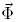* fits and mRNA provide only a 7% to 15% increase in explanatory power over the *without 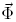* ROC SEMPPR predictions of *ϕ*. Similarly, correlation between *ϕ* estimates for ROC SEMPPR’s *with 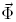* fits and RPF reads provide a 6% to 12% increase in explanatory power over its *without 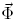* predictions of *ϕ*.

**Figure 5:**
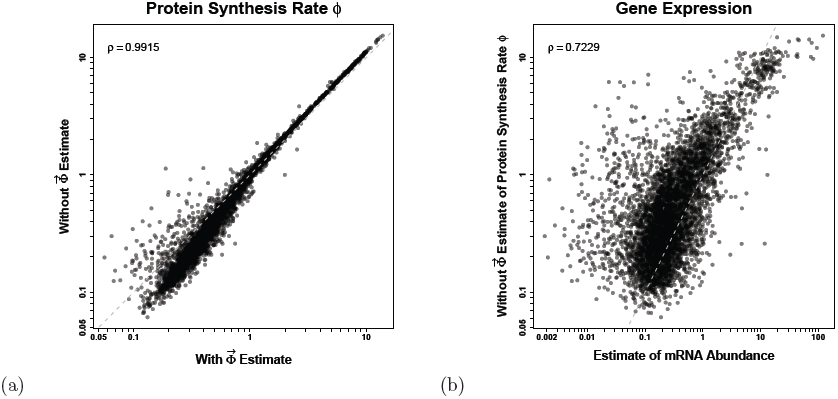
Evaluation of predicted gene expression levels between models and empirical measurements from Yassour *et al*. (2009). (a) Comparison of *with* and *without 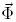* ROC SEMPPR estimates of protein synthesis rates,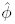. The units for *ϕ* are protein/*t* and time *t* is scaled such that the prior for *ϕ* satisfies *E*(*ϕ*) = 1. Note the very strong correlation between the *with* and *without 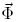*estimates of *ϕ* for the high expression genes. (b) Comparison of *without 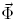* estimates of *ϕ* and empirical measurements of mRNA abundances, 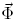. The empirical mRNA abundance measurements, [mRNA], are being used here as a proxy for protein synthesis rates, i.e. [mRNA] protein/*t*. The measurements are scaled such that the mean [mRNA] value is 1. Pearson correlation coefficients *ρ* are given and the dashed gray line indicates 1:1 line.

**(Approximate Location of Figure 5)**

### Changes in CUB with Protein Synthesis Rate

As first shown in Shah and Gilchrist (2011a), the relationship between codon usage and protein synthesis rate *ϕ* can range from simple and monotonic to complex. Figure 6 illustrates how codon usage changes across approximately 2 orders of magnitudes of 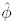 for each of the *n*_aa_ = 19 multicodon amino acids. Both ROC SEMPPR’s *with* and *without 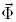* model fits accurately predict how CUB changes with protein synthesis rates (Figure 6). Indeed, the predicted changes in CUB between the *with* and *without 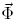*ROC SEMPPR model fits are almost indistinguishable from one another, reflecting the strong agreement between their estimates of Δ*M* and Δ*η* across models as discussed above.

Changes in codon frequency with *ϕ* are the result of a subtle interplay between natural selection for reducing *η* and mutation bias (Figure 6). The simplest cases involve two codon amino acids where the same codon is favored both by selection and mutation bias, i.e. Cys, Glu, and Ser_2_. In these three cases, the selectively and mutationally favored codon 1 is used preferentially across all protein synthesis rates and the frequency of the preferred codon increases monotonically with *ϕ*. The next simplest cases involve two codon amino acids where codon 1 is favored by selection and codon 2 is favored by mutation bias, e.g. Asp, Asn, and Phe. In these cases, the mutationally favored codon 2 is used preferentially at low *ϕ* values and the selectively favored codon 1 is used preferentially used in genes at high *ϕ* values. Nevertheless, as before the codon frequency changes monotonically with *ϕ*.

**Figure 6:**
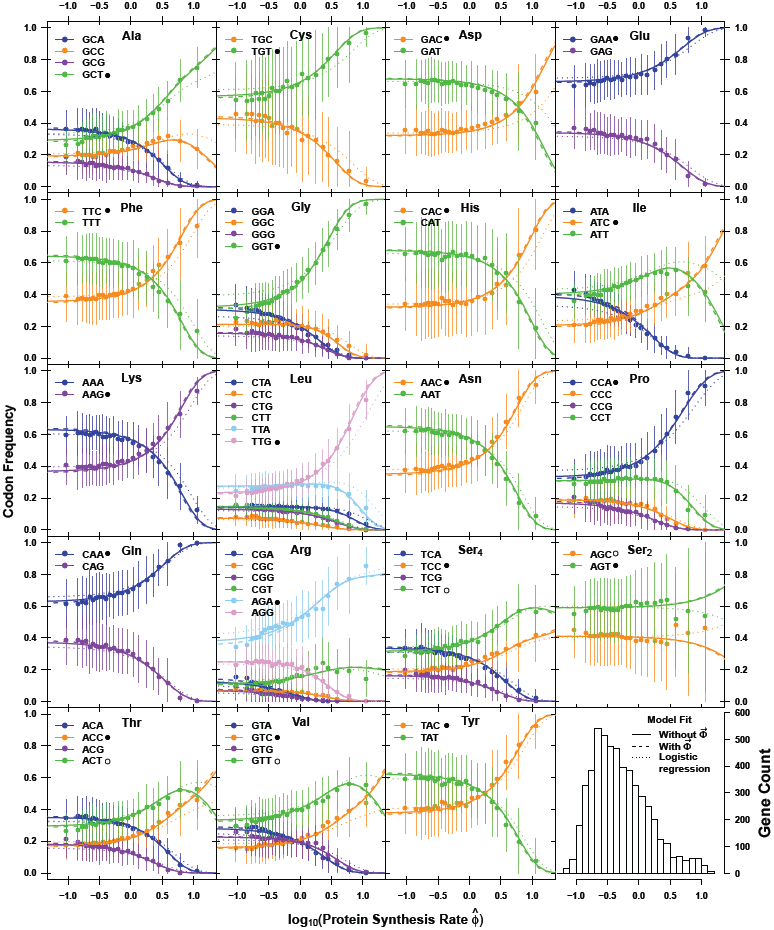
Model predictions and observed codon usage frequencies as a function of estimated protein synthesis rate *ϕ* for the *S. cerevisiae* S288c genome. The units for *ϕ* are protein/*t* and time *t* is scaled such that the prior for *ϕ* satisfies *E*(*ϕ*) = 1. Each amino acid is represented by a separate subplot. Solid, dashed, and dotted lines represent the *without 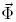*, *with 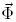* ROC SEMPPR model fits, and a simple logistic regression approach where the estimation error in 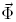 is ignored, respectively. None of the parameter estimates’ 95% Credibility Intervals overlap with 0 except Δ_*ηCGT,AGA*_. Genes are binned by their expression levels with solid dots indicating the mean codon frequency of the genes in the respective bin. Error bars indicate the standard deviation in codon frequency across genes within a bin. For each amino acid, the codon favored by natural selection for reducing translational inefficiency is indicated by a •. The four indicate codons that have been previously identified as ‘optimal’ but our ROC SEMPPR model fits indicate these codons actually are the second most efficient codons. A histogram of the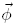values is presented in the lower right corner. Estimates of protein synthesis rates 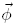 are based on the *with 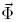* ROC SEMPPR model fits, thus representing our best estimate of their values.

**(Approximate Location of Figure 6)**

More complex, non-monotonic changes in codon frequencies can occur in amino acids that use three or more codons. For example, the Ile codon ATC has the lowest translational inefficiency Δ*η* and, therefore, is the most favored codon by natural selection while ATT has the second lowest translational inefficiency. As a result, both codons initially increase in frequency with increasing *ϕ* at the expense of the most inefficient codon ATA. However, once the frequency of ATA approaches 0, selection for ATC begins driving the frequency of ATT down. These non-monotonic changes in codon frequency is most notable in Ala, Ile, Thr, and Val. Examining the derivative of 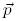 with respect to *ϕ* indicates that if 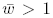, a given codon *i* will increase in frequency with *ϕ*, if 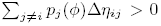 i.e. if the sum of the derivatives of the selective advantage of codon *i* over the other codons is positive. For the reference codon 1 where, by definition, Δ*η*_*i,*1_ *≥* 0, we see that this inequality *always* holds. This criteria can only be met by the non-reference codon in amino acids with more than two synonyms and when there are other non-reference codons with lower fitnesses at appreciable probabilities. In the *S. cerevisiae* S288c genome, these conditions can occur when the codon most favored by natural selection is strongly disfavored by mutation. Although this non-linear quality of multinomial logistic regression is well known among statisticians, the fact that non-optimal codons other than the choice most favored by selection can increase with production rate has not been widely recognized by biologists.

If we ignore the noise in the *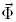* data, our *with 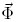* model fitting simplifies to the standard logistic regression model applied in Shah and Gilchrist (2011a). This simplification results in a slight distortion of Δ*M* estimates and a general attenuation of our estimates of Δ*η* (Wallace *et al*., 2013). The effect of this attenuation can be seen in Figure 6 where the changes in CUB predicted from the standard logistic regression model fit lag behind the predicted changes when either the error in 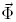 is accounted for or the *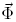* data is not used. In the case of Ser_2_ controlling for error leads to a change in the codon identified as being favored by natural selection. While Shah and Gilchrist (2011a) predicted codon AGC would be favored by selection over AGT, both of ROC SEMPPR’s *with* and *without 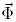* fits predict the opposite. Although, this switch in order is ‘significant’ in that the 95% CI for Δ*ηAGT,AGC* is *<* 0, the amino acid Ser_2_ is used at very low frequency in high expression genes and its 97.5% CI boundary lies very close to 0. (The upper boundary lies at 0.00387 and 0.000634 for the *with* and *without 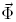*ROC SEMPPR fits, respectively.) As a result, this discrepancy is not strongly supported and warrants further investigation.

In summary, for genes with protein synthesis rates substantially lower than the average, i.e. 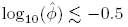, codon usage is largely determined by mutation bias terms Δ*M*. For about half of the amino acids (e.g. Cys, Lys, and Pro), in genes with protein synthesis rates 10 or more times greater than average, i.e. 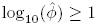, codon usage is largely determined by selection for the codon with the smallest translational inefficiency Δ*η*. This result is largely consistent with the frequent assumption that in the set of genes with the highest expression levels the most translationally efficient codon dominates. However, for the amino acids (e.g. Ala, Ile, and Arg) selection for reducing *η* in high expression genes is substantially tempered by the force of mutation bias.

### Estimating Selection on Synonymous Codon Usage

The assumptions of the ROC SEMPPR model imply that the codon specific translational inefficiencies are independent of codon position within a sequence. As a result, the relative strength of purifying selection on synonymous codon *j* in comparison to codon *i* in a gene with an average protein synthesis rate *ϕ* is,

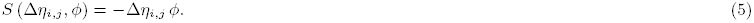

We remind the reader that Δ*η* includes the effective population size, *N*_*e*_, in its definition. As a result, our selection coefficients *S* are measured relative to the strength of genetic drift, 1*/N_e_*, as is commonly done. The distribution of *S* across all genes for each alternative to an amino acid’s reference codon are illustrated in Figure 7 and summarized in Table 1. Tables with genome wide gene and codon specific estimates of *S* can found in the Supporting Materials. Recall that *S* is scaled by *ϕ* and that the distribution of *ϕ* values across genes appears to follow a heavy tailed distribution. As a result even though, by definition, the average value of *ϕ* is 1, the large majority of genes have *ϕ* values less than 1. As a result, although purifying selection on synonymous codons is universal, its selection coefficients are usually quite small (i.e. *> −*0.5). Nevertheless, because our framework utilizes information on CUB held across genes, we can clearly detect the signature of selection at the genome level, specifically in the form of Δ*η* values whose posterior credibility intervals differ from 0, while other approaches might fail.

**Table 1:** Summary statistics for gene specific selection coefficients on synonymous codon usage *S* = *−*Δ*η ϕ* from the *without 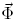* ROC SEMPPR model fit to the *S. cerevisiae* genome. The selection coefficient *S* was calculated relative to the most translationally efficient codon for a given amino acid on a gene by gene basis.

| AA | Codon | $\bar{S}$ | $\text{var}(S)$ | Quantile | | | | | | |
| --- | --- | --- | --- | --- | --- | --- | --- | --- | --- | --- |
|  |  |  |  | 1% | 5% | 25% | 50% | 75% | 95% | 99% |
| A | GCA | -0.4743 | 0.6171 | -4.3994 | -1.7817 | -0.4715 | -0.2096 | -0.1135 | -0.0620 | -0.0466 |
| A | GCC | -0.0648 | 0.0115 | -0.6014 | -0.2436 | -0.0645 | -0.0287 | -0.0155 | -0.0085 | -0.0064 |
| A | GCG | -0.6420 | 1.1306 | -5.9551 | -2.4117 | -0.6382 | -0.2837 | -0.1536 | -0.0839 | -0.0631 |
| C | TGC | -0.3777 | 0.3913 | -3.5032 | -1.4187 | -0.3754 | -0.1669 | -0.0903 | -0.0493 | -0.0371 |
| D | GAT | -0.1201 | 0.0396 | -1.1142 | -0.4512 | -0.1194 | -0.0531 | -0.0287 | -0.0157 | -0.0118 |
| E | GAG | -0.2527 | 0.1752 | -2.3440 | -0.9493 | -0.2512 | -0.1117 | -0.0605 | -0.0330 | -0.0248 |
| F | TTT | -0.2741 | 0.2061 | -2.5425 | -1.0297 | -0.2725 | -0.1211 | -0.0656 | -0.0358 | -0.0269 |
| G | GGA | -0.8286 | 1.8834 | -7.6861 | -3.1127 | -0.8237 | -0.3662 | -0.1982 | -0.1083 | -0.0815 |
| G | GGC | -0.4436 | 0.5398 | -4.1149 | -1.6664 | -0.4410 | -0.1960 | -0.1061 | -0.0580 | -0.0436 |
| G | GGG | -0.7052 | 1.3641 | -6.5412 | -2.6490 | -0.7010 | -0.3116 | -0.1687 | -0.0921 | -0.0693 |
| H | CAT | -0.1966 | 0.1060 | -1.8233 | -0.7384 | -0.1954 | -0.0869 | -0.0470 | -0.0257 | -0.0193 |
| I | ATA | -0.8874 | 2.1604 | -8.2318 | -3.3337 | -0.8822 | -0.3922 | -0.2123 | -0.1159 | -0.0872 |
| I | ATT | -0.0906 | 0.0225 | -0.8403 | -0.3403 | -0.0901 | -0.0400 | -0.0217 | -0.0118 | -0.0089 |
| K | AAA | -0.2661 | 0.1943 | -2.4686 | -0.9997 | -0.2646 | -0.1176 | -0.0637 | -0.0348 | -0.0262 |
| L | CTA | -0.2636 | 0.1906 | -2.4452 | -0.9902 | -0.2620 | -0.1165 | -0.0631 | -0.0344 | -0.0259 |
| L | CTC | -0.7185 | 1.4162 | -6.6650 | -2.6991 | -0.7143 | -0.3175 | -0.1719 | -0.0939 | -0.0706 |
| L | CTG | -0.4662 | 0.5961 | -4.3242 | -1.7512 | -0.4634 | -0.2060 | -0.1115 | -0.0609 | -0.0458 |
| L | CTT | -0.4601 | 0.5807 | -4.2679 | -1.7284 | -0.4574 | -0.2033 | -0.1101 | -0.0601 | -0.0452 |
| L | TTA | -0.2128 | 0.1242 | -1.9741 | -0.7995 | -0.2116 | -0.0941 | -0.0509 | -0.0278 | -0.0209 |
| N | AAT | -0.3164 | 0.2746 | -2.9347 | -1.1885 | -0.3145 | -0.1398 | -0.0757 | -0.0413 | -0.0311 |
| P | CCC | -0.5061 | 0.7027 | -4.6948 | -1.9013 | -0.5031 | -0.2237 | -0.1211 | -0.0661 | -0.0498 |
| P | CCG | -0.8002 | 1.7567 | -7.4230 | -3.0061 | -0.7955 | -0.3537 | -0.1914 | -0.1046 | -0.0787 |
| P | CCT | -0.2319 | 0.1476 | -2.1513 | -0.8712 | -0.2306 | -0.1025 | -0.0555 | -0.0303 | -0.0228 |
| Q | CAG | -0.3840 | 0.4046 | -3.5624 | -1.4427 | -0.3818 | -0.1697 | -0.0919 | -0.0502 | -0.0378 |
| R | AGG | -0.5378 | 0.7935 | -4.9890 | -2.0204 | -0.5347 | -0.2377 | -0.1287 | -0.0703 | -0.0529 |
| R | CGA | -1.7330 | 8.2391 | -16.0757 | -6.5103 | -1.7228 | -0.7659 | -0.4146 | -0.2264 | -0.1704 |
| R | CGC | -0.5568 | 0.8504 | -5.1646 | -2.0915 | -0.5535 | -0.2461 | -0.1332 | -0.0727 | -0.0547 |
| R | CGG | -1.5092 | 6.2486 | -13.9999 | -5.6696 | -1.5003 | -0.6670 | -0.3611 | -0.1972 | -0.1484 |
| R | CGT | -0.0080 | 0.0002 | -0.0744 | -0.0301 | -0.0080 | -0.0035 | -0.0019 | -0.0010 | -0.0008 |
| S | TCA | -0.3942 | 0.4264 | -3.6571 | -1.4810 | -0.3919 | -0.1742 | -0.0943 | -0.0515 | -0.0388 |
| S | TCG | -0.4927 | 0.6660 | -4.5705 | -1.8509 | -0.4898 | -0.2178 | -0.1179 | -0.0644 | -0.0484 |
| S | TCT | -0.0121 | 0.0004 | -0.1120 | -0.0454 | -0.0120 | -0.0053 | -0.0029 | -0.0016 | -0.0012 |
| T | ACA | -0.4278 | 0.5020 | -3.9679 | -1.6069 | -0.4252 | -0.1890 | -0.1023 | -0.0559 | -0.0421 |
| T | ACG | -0.6503 | 1.1603 | -6.0327 | -2.4431 | -0.6465 | -0.2874 | -0.1556 | -0.0850 | -0.0639 |
| T | ACT | -0.0482 | 0.0064 | -0.4468 | -0.1810 | -0.0479 | -0.0213 | -0.0115 | -0.0063 | -0.0047 |
| V | GTA | -0.6310 | 1.0922 | -5.8529 | -2.3703 | -0.6272 | -0.2789 | -0.1509 | -0.0824 | -0.0620 |
| V | GTG | -0.4308 | 0.5092 | -3.9966 | -1.6185 | -0.4283 | -0.1904 | -0.1031 | -0.0563 | -0.0424 |
| V | GTT | -0.0570 | 0.0089 | -0.5288 | -0.2142 | -0.0567 | -0.0252 | -0.0136 | -0.0074 | -0.0056 |
| Y | TAT | -0.2857 | 0.2239 | -2.6501 | -1.0732 | -0.2840 | -0.1263 | -0.0683 | -0.0373 | -0.0281 |
| Z | AGC | -0.0248 | 0.0017 | -0.2297 | -0.0930 | -0.0246 | -0.0109 | -0.0059 | -0.0032 | -0.0024 |

**Figure 7:**
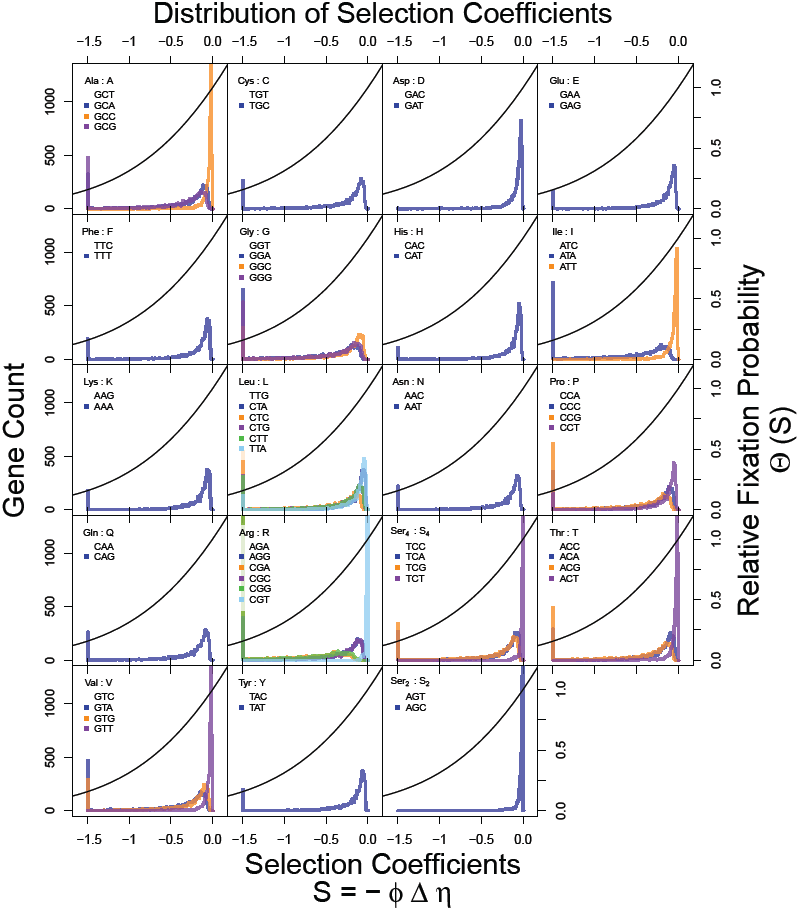
Distribution of gene specific selection coefficients on synonymous codon usage *S* = *−*Δ*η ϕ* from the *without 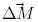* model fit to the *S. cerevisiae* genome. Selection coefficient *S* were calculated on a gene by gene basis and relative to the most translationally efficient codon for a given amino acid (which is the codon listed first in the legend). The reference codon, which is most favored by selection and for which, by definition, *S* = 0, is listed first within the legend of each panel. Genes with *S* 2 were combined together into a single bin. For reference, the fixation probability of a codon relative to a pure drift process, Θ (*S*) = 2*S/*(1 *−* exp [*−*2*S*]), are also plotted (– line). Summary statistics can be found in Table 1.

**(Approximate Location of Figure 7)**

**(Approximate Location of Table 1)**

Figure 8 compares our *without 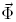*ROC SEMPPR based estimates of *S* to those estimated using the FMutSel phylogenetic model of Yang and Nielsen (2008) using PAML (Yang, 2007) for the 106 genes in the Rokas *et al*. (2003) dataset. Overall we observe reasonable qualitative agreement between the two models with the majority of codon specific predictions having correlation coefficients *ρ >* 0.3. Unfortunately, while PAML provides maximum likelihood point estimates of parameters, it does not provide any confidence intervals for these parameters. Given the large number of parameters (*>* 60) estimated from each coding sequence by FMutSel, the confidence intervals for each parameter is likely to be large and, hence, could explain much of the variation we observe between ROC SEMPPR and FMutSel parameter estimates. Nonetheless, for 85% of the codons examined (34/40), we observe is a significant (*p <* 0.05) and positive linear relationship between the ROC SEMPPR and the FMutSel estimates of *S* (see Table S11). Of the remaining 6 codons, half exhibit a positive, but non-significant relationship between ROC SEMPPR and FMutSel’s estimates of *S*, while the other half exhibit a negative, but again non-significant, relationship between estimates of *S*. Thus for 92% of the codons, both the ROC SEMPPR and FMutSel estimates of *S* agree qualitatively.

The three exceptions to this qualitative agreement are codons CGT (Arg), TCT (Ser4), and ACT (Thr) and it is worth noting two points. First, the central 95% CI for CGT (Arg) overlaps with 0 in both the *with* and *without 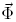* ROC SEMPPR model fits. Second, the Ser_4_ codon TCT and Thr codon ACT are two of the four codons that ROC SEMPPR indicates have been misidentified as ‘optimal’ codons in the past. Relative to the ROC SEMPPR reference codons, TCT and ACT have small Δ*η* values, *˜* 0.01, and *∼* 0.05 respectively, and large Δ*M* values, *~ −*0.5 for both. Thus, it appears in these last two cases the FMutSel model is misattributing the CUB towards these codons to selection rather than mutation (see Figure 6).

**Figure 8:**
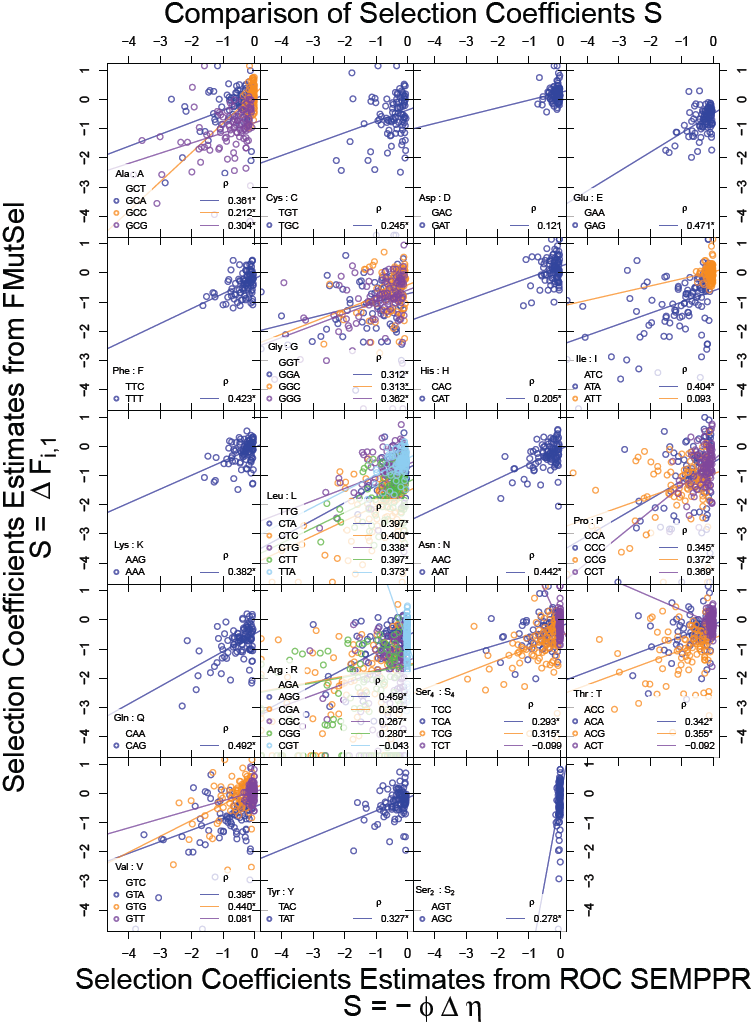
Comparison of gene specific selection coefficients on synonymous codon usage *S* = *−*Δ*η ϕ* from the *without* 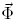 model fit to the *S. cerevisiae* genome and those from fitting the FMutSel model from Yang and Nielsen (2008) for 106 yeast genes used in Rokas *et al*. (2003) as estimated by Kubatko *et al*. (view) For more details see the main text. Selection coefficient *S* were calculated on a gene by gene basis and relative to the most translationally efficient codon for a given amino acid (which is the codon listed first in the legend). Lines indicate linear regression line best fit and the corresponding correlation coefficients are listed as well with a *\** indicating model fits with *p <* 0.05. Under the FMutSel model, monomorphic sites across species can lead to estimates of *S* = *−∞*, these observations are plotted on the x-axis.

**(Approximate Location of Figure 8)**

## Discussion

Recent advances in technology have led a remarkable and continuing decrease in the cost of genome sequencing. What is now needed are robust models and computational tools that allow researchers to access the information encoded within these genomes. Several models have been proposed that estimate selection coefficients of all 61 sense codons either on a whole gene basis or on a site-by-site basis (Tamuri *et al*., 2012; Rodrigue *et al*., 2010; Yang and Nielsen, 2008). While important advances, these models fail to leverage information on CUB encoded across genes. In contrast, ROC SEMPPR estimates selection coefficients and other key parameters by assuming a common directionality of selection on CUB, but where the strength of selection varies with protein synthesis rate.

As a result, ROC SEMPPR provides a modeling framework which can quickly extract information on codon specific translational inefficiencies Δ*η*, mutation biases Δ*M*, and gene specific estimates of protein synthesis rates 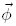, using only genome wide patterns of CUB. This ability stems from the hypotheses that the intergenic variation in patterns of CUB observed within a genome reflect a lineage’s evolutionary responses to selection against inefficient protein translation as well as mutation bias. Our results clearly show that these CUB patterns contain remarkably large amounts of useful quantitative information and the use of carefully constructed, mechanistically driven mathematical models can greatly improve our ability to access and interpret this information. Indeed, we find that for *S. cerevisiae* ROC SEMPPR’s *without 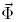* estimates of Δ*M*, Δ*η*, and 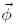 values match almost exactly with the *with 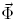* estimates of these parameters. By removing the need for gene expression data 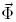 and, instead, providing reliable predictions of their average protein synthesis rates 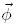, the methods developed here should be especially helpful for molecular-, systems-, and micro-biologists for whom genomic sequence data are both abundant and inexpensive to obtain. For example, the protein translation rates we estimate 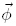 should contain useful information about the physiology and ecology of the organism. Indeed, for the large number of sequenced micro-organisms that cannot be easily cultured in the laboratory, their genome sequence may become the primary source of information about their biology for the near future.

Of course ROC SEMPPR may not work for all organisms. For example, some organisms may evolve under *N*_*e*_ values too small for adaptation in CUB to occur. Under these conditions, our method should fail to confidently identify the selectively preferred codon (i.e. our credibility intervals for our Δ*η* parameters will overlap with 0). However, because our estimates of Δ*η* are based on the analysis of the entire genome simultaneously rather than the combination of independent assessments of individual genes, our method may be able to detect the signature of selection on CUB in organisms where it previously went undetected. Alternatively, there may be organisms where *N*_*e*_ is so large that, as a result, there is not enough variation in CUB to reliably estimate our parameters. Assuming we retain our flat priors on Δ*η*, in these cases we expect our estimates of Δ*η* for the most selectively favored codons to continually increase in magnitude rather than eventually stabilizing. Such behavior reflects a lack of information in the data rather than a flaw in our model and has been observed in other approaches, such as those using inter-and intra-specific variation (Yang and Nielsen (2008) and Lawrie *et al*. (2013), respectively). ROC SEMPPR may also fail to work with organisms whose adaptation in CUB are driven by more complex or less consistent selective forces. If these forces are uncorrelated across amino acids within a gene or varied greatly with position within a gene, then our method should not be able to confidently identify the selectively preferred codons, similar to the case with of organisms with small *N*_*e*_.

While direct, codon specific estimates of Δ*M* and Δ*η* do not exist, data from mutation accumulation lines and tRNA copy number can be used as proxies. Reassuringly, we observe strong and consistent agreement between ROC SEMPPR’s parameter estimates and these proxies. In addition, when comparing ROC SEMPPR’s estimates of *S* to the FMutSel model we observe general agreement between our estimates with the three key exceptions likely due to relatively small differences in translational inefficiencies between these synonymous codons and their most efficient alternative and strong mutation bias against the most efficient, which is misinterpreted by FMutSel as selection. In contrast, the selective values *S* on synonymous codon usage we estimate using ROC SEMPPR are substantially smaller than those estimated by Lawrie *et al*. (2013) based on intra-specific variation in four fold degenerate sites for *Drosophila melanogaster*. While we have no immediate explaination for these differences we do note that Lawrie *et al*. (2013) acknowledge that the high *S* values they estimate are the exception rather than the rule for population genetic studies, including those looking at non-synonymous substitutions.

Because of ROC SEMPPR’s derivation from population genetics, it should be possible to take any observed intra-specific variation into account by expanding our codon counts likelihood function in equation (8) to be calculated across the polymorphic alleles in proportion to their frequencies. Further, given that the directionality of selection in ROC SEMPPR is estimated using information from across the genome, our ability to detect site specific violations of the model should be much greater than when analyzing the CUB of each gene separately. Expanding ROC SEMPPR to utilize inter-specific variation, however, is more complex and will require expanding the model to include the effects of non-synonymous substitutions and phylogenetic history.

For organisms that can be cultured in the laboratory, researchers can utilize experimental techniques to measure mRNA, ribosome profiles, and protein abundances. Even though impressive gains have been made in our ability to measure these quantities at a genome scale, abundance data still have limitations. For example, mRNA abundance measurements have been shown to vary substantially between labs using the same strain and the same general conditions (Wallace *et al*., 2013). Indeed, our posterior mean estimate of the error in mRNA abundance measurement (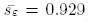) indicates that the error in a given measurement ranges over an order of magnitude. In terms of protein abundance measurements, most proteomic studies have difficulty quantifying membrane bound proteins [Durr *et al*. (2004); Babu *et al*. (2012); Chen *et al*. (2013)]. Furthermore, both transcriptomic and proteomic measurements are, by their very nature, restricted to the specific growth conditions used. Unfortunately, the frequency with which organisms outside of the lab encounter such conditions is generally unknown. This is particularly important for understanding a pathogenic organism, where expression of genes involved in its persistence and spread are highly dependent on their hosts and are difficult to mimic *in vitro*.

The predictions of protein synthesis rates 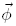 generated by ROC SEMPPR contain independent and complementary information to that found in mRNA or protein abundance measurements. As a result, this information can be used on its own or in combination with other measures of gene expression. For example, our work provides estimates of protein production based on the average environment that an organism’s lineage has experienced. These estimates of average gene expression can be used to further contextualize gene expression measurements in different environments. For example, comparing the *ϕ* values for proteins involved in different, environment specific pathways should give researchers an understanding of the relative importance these environments in the lineage’s evolutionary history. At a finer scale, gene-specific incongruences between mRNA abundance measurements and *ϕ* estimates may indicate genes undergoing extensive post-transcriptional regulation, a hypothesis that can be evaluated experimentally.

The fact that the additional information provided by the 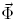 data from Yassour *et al*. (2009) leads to a relatively small increase in the quality of our predictions of *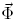*data from other labs may seem surprising. However, we believe this behavior indicates that the information in *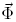* about gene specific protein synthesis rates is largely redundant with the information held within the CUB patterns within a gene and across a genome.

Of course a skeptic might proffer a different interpretation, i.e. the model is somehow ignoring or insensitive to the information in 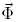. We, however, believe this is not the case for the following reasons. First, ROC SEMPPR was carefully formulated to combine the information from independent *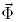* measurements and the CUB of each gene in a straightforward and logical manner (See Supporting Materials: Fitting of Model to Genomic Data and Noisy Measurements and Equation (S1) in particular). Instead of *a priori* assuming one source of information is better than the other, ROC SEMPPR actually evaluates the relative quality of each source of information in explaining the observed bias in codon usage for a gene across the *n*aa amino acids. Next, given the fact that the 95% Posterior Credibility Intervals for Δ*η* differ for at least one pair of codons for each amino acid indicates that the information held within the CUB patterns is reliable. In contrast, ROC SEMPPR’s estimate of the error in *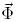* indicates that the empirical measurements *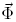* are noisy, consistent with the findings from other studies. For example, Wallace *et al*. (2013) looked at the the correlation in *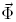* measurements between independent labs and found non-trivial disagreement in their values. Finally, and perhaps most convincingly, the *without* 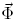 version of ROC SEMPPR treats the *ϕ* values as missing values and is able to predict their values to a similar level of accuracy observed between empirical measurements from different laboratories and using different platforms (Supporting Figures S4 and S5).

Accessing information on 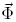 using a mechanistic, model based approach as developed here has additional, distinct advantages over more *ad-hoc* approaches frequently used by other researchers. Quantifying selection on synonymous codons is important for phylogenetic inference. Classical codon substitution models of protein evolution typically assume that synonymous codons of an amino acid are selectively neutral. In contrast, our estimates of codon-specific translation inefficiencies Δ*η* and expression levels 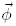 provide an independent measure of selection on synonymous codons from a single genome. By incorporating these measures in codon substitution models, researchers would be able to measure selection on non-synonymous changes either within a gene or on a site-by-site basis.

In addition, current measures to identify the selective regime in which a gene evolves, e.g. positive, negative or nearly-neutral, are based on estimating the number of non-synonymous to synonymous changes (dN/dS) (Li *et al*., 1985; Nei and Gojobori, 1986; Yang and Nielsen, 2000) or polymorphism data (McDonald and Kreitman, 1991). These tests generally assume that synonymous changes are neutral. However, (Spielman and Wilke, 2015) have recently shown, ignoring selection on synonymous changes can lead to a false positive signal of a gene evolving in response to diversifying selection. By using our codon-specific estimates of translation inefficiencies, researchers will now be able to explicitly account for biases in estimates of dS due to selection on synonymous changes (Spielman and Wilke, 2015).

Estimates of codon-specific translation inefficiencies are also important for practical applications such as codon-optimization algorithms that are used to increase heterologous gene expression, for e.g. insulin expression in *E. coli*. When heterologous genes are expressed in a particular model organism such as *E. coli* or *S. cerevisiae*, their codon usage is ‘optimized’ by assuming that the most frequently used codon in a set of highly expressed genes is the optimal one. This approach implicitly assumes that natural selection against translational inefficiencies overwhelms any mutation bias. In several amino acids that use more than two synonymous codons, e.g. Ser4, Thr and Val, genes with highest expression are more often encoded by the mutationally favored, second-best codon rather than the mutationally disfavored ‘optimal’ codon. As a result, relying on the codon usage of highly expressed genes appears to be overly simplistic in the case of the *S. cerevisiae* genome and, if our inferences are correct, has led to misidentification of the ‘optimal’ codon.

In addition to codon-specific translation inefficiencies Δ*η*, we also estimate codon-specific mutation biases Δ*M*. We find that the direction of mutation biases between synonymous codons is consistent across all amino acids and in the same direction as genomic AT content. However, as we documented in Shah and Gilchrist (2011a), Δ*M* for similar sets of nucleotides differ significantly between amino acids. For instance, in the case of two-codon amino acids with C-T wobble, we find that Δ*MN N C,N N T* ranges from 0.27 to 0.75. For genes with low expression levels (i.e. *ϕ <* 1), this corresponds to ratios of T-ending codons to C-ending codons between amino acids ranging from 1.3 to 2.1. One possible explanation for this wider than expected range of mutation biases could be context-dependence of mutation rates. Recent high-throughput mutation accumulation experiments in yeast support this idea, estimating that the mutation rate at a particular nucleotide depends on the context of surrounding nucleotides: the **C** nucleotide in the context of C**C**G has several fold higher mutation rate than in the context of C**C**T (Zhu *et al*., 2014).

Despite the numerous advances outlined above, our work is not without its limitations. One important limitation stems from our assumption that codons contribute to the cost-benefit ratio of protein translation in an additive manner. While this assumption is consistent with certain costs of protein translation, such as ribosome pausing, it ignores many others selective forces potentially shaping the evolution of CUB. For example, the cost of nonsense errors, i.e. premature termination events, are generally expected to increase with codon position along an ORF and, thus, lead to a non-additive contribution of a given codon to the cost-benefit ratio *η* (Gilchrist *et al*., 2009). Similarly, if one assumes that the main effect of missense errors is to reduce the functionality of the protein produced, then the cost of these errors is expected to depend greatly on specific details such as the structural and functional role of the amino acid at which the error occurs and the physiochemical differences between the correct and the erroneously incorporated amino acids. Finally, the pausing time at a codon is also influenced by several factors such as downstream mRNA folding (Yang *et al*., 2014), presence of polybasic stretches (Brandman *et al*., 2012) as well as co-translational folding of the growing polypeptide (Thanaraj and Argos, 1996; Pechmann and Frydman, 2013). While the contributions of these factors to ribosomal pausing times are often idiosyncratic and vary widely between genes, they can all influence the cost-benefit ratio *η*. The situation becomes even more complex and non-linear when considering how nonsense and missense errors along with various factors influencing pausing time costs combine to affect *η*. In all of these situations, the nonlinear mapping between a codon sequence and *η* makes direct evaluation of the likelihood function difficult. In such situations alternative, approximate methods and simulation techniques, such as those developed by (Murray *et al*., 2006), will become necessary. Expanding our approach to include these additional selective forces should allow us to quantitatively evaluate the separate contributions of ribosome pausing time, nonsense errors, and missense errors have made to the evolution of CUB for a given species. Doing so will allow us to address the long held goal in molecular and evolutionary biology of accurately quantifying the factors contributing to the evolution of CUB within a coding sequence and across a genome.

## Methods and Materials

### Modeling Natural Selection on Synonymous Codons

Following the notation and framework introduced in Gilchrist (2007) and Shah and Gilchrist (2011a), we assume that for each gene, the organism has a target, average protein synthesis rate *ϕ*. Protein synthesis rates have units of 1/time; for convenience and ease of interpretation, we define our time units such that the average or expected protein synthesis rate across the genome is one, i.e. *E*(*ϕ*) = 1. The cost-benefit ratio 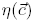 represents the expected cost, in ATPs, to produce one functional protein from the coding sequence 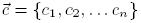 where *c*_*i*_ represents the codon used at position *i* in a protein of length *n*. In its most general form, 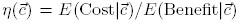, where *E*(Cost) is the expected direct and indirect energetic costs incurred by a cell when a ribosome initiates translation of a transcript containing 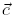. Similarly, *E*(Benefit|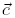) is the expected benefit, relative to a complete and error free protein, received by a cell when a ribosome initiates translation of a transcript containing 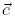. By definition, in the absence of translation errors, ribosomes will only produce complete and error free proteins, i.e. for ROC SEMPPER *E*(Benefit) = 1. Thus any differences in *η* are the result of differences in *E*(Cost) between alternative 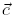’s and *E*(*Cost*) simplifies to 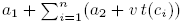 where *a*_1_ is the direct and indirect cost of translation initiation, *a*_2_ is the direct cost of peptide elongation (4 ATPs/amino acid), *t*(*c*_*i*_) is the average pausing time a ribosome takes to translate codon *ci*, and *v* scales this indirect cost of ribosome pausing from units of time to ATPs. Based on these definitions, 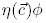 represents the average energy flux an organism must expend to meet its target production rate for a given protein. If we assume that every ATP/time spent leads to a small, proportional reduction in genotype fitness *q*, then the fitness of a given genotype is,

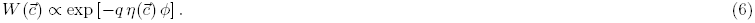

In the simplest scenarios, such as when there is selection to minimize ribosome pausing during protein synthesis, a synonymous codon *i* makes an additive, position independent contribution to *η*. In this scenario, the evolution of the codons in 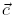is independent between positions. As a result, the information held within 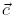can be summarized by the number of times each synonymous codon is used within 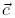. Given these assumptions, within the ORF of a given gene the stationary probability of observing a set of codon counts 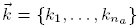 for a given amino acid with *n*_*a*_ synonymous codons within 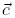 will follow a multinomial distribution with the probability vector *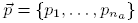*. Here, for *i* = 1*, …, na*,

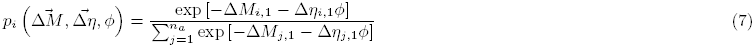

where Δ*M*_*i,*1_ is a measure of codon specific mutation bias and Δ*η*_*i,*1_ is a measure of translational inefficiency. Specifically, Δ*M_i,1_* = ln (*p*_1_*/p_i_*) |_*ϕ*=0_, that is the natural logarithm of the ratio of the frequencies of synonymous codon 1 to *i* in the absence of natural selection. Following the detailed balance assumptions in our population genetics model, in the specific cases where codons *i* and 1 can mutate directly between each other, Δ*M_i,1_* is also equal to the log of the ratio of the mutation rates between the two codons (Sella and Hirsh, 2005a; Shah and Gilchrist, 2011a; Wallace *et al*., 2013). Following Sella and Hirsh (2005a), for *N*_*e*_≫ 1, for both a haploid and diploid Fisher-Wright populations, we scale the differences in the contribution two synonymous codons make to *η* relative to genetic drift, i.e. Δ*η_i,j_* = 2*Ne* (*η_i_ − η_j_*). Because the reference codon 1 is determined by pausing time values, Δ*M_i,1_* values can be both negative and positive, unlike Δ*η_i,1_*.

### Fitting the Model to Genomic Data

Our main goal is to estimate codon specific differences in mutation bias,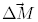, translational inefficiencies, 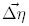 and protein synthesis rates for all genes, 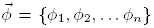 from the information encoded in the codon usage patterns found across a genome. To test our approach we used the *S. cerevisiae* S288c genome file orf_coding.fasta.gz which was posted on 03 February 2011 by Saccharomyces Genome Database http://www.yeastgenome.org/ (Engel *et al*., 2014)). This data contains 5,887 genes and consists of the ORFs for all “Verified” and “Uncharacterized” genes as well as any transposable elements. To fit the *with 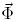* model we used RNA-seq derived mRNA abundance measurements from Yassour *et al*. (2009). We combined the abundance measures from the four samples, YPD0.1, YPD0.2, YPD15.1, and YPD15.2, taken during log growth phase and used the geometric mean of these values as a proxy for relative protein synthesis rates *ϕ′*. As is commonly done by empiricists, we rescaled our *ϕ′* values such that they summed to 15,000. Because our *with 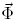* model fits estimate the scaling term, exp(*A*F), the only effect of this rescaling is on our estimate of *A*F. To reduce noise in the 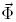 data, we only used genes with at least three non-zero measurements. The intersection of 5,887 DNA ORF sequences and 6,303 mRNA abundance measurements produced 5,346 ORF’s in common to both datasets. These 5,346 genes made up the final dataset used for ROC SEMPPR’s *with* and *without 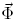* model fits.

Using an MCMC approach we sample from the posterior distribution, according to the equation

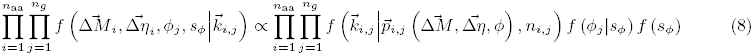

where the likelihood of the codon counts,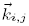, are naturally modeled as a multinomial distribution (Multinom) for the amino acid *i* in the ORF of gene *j* as defined in Equation (7), 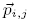is an inverse multinomial logit function (mlogit^*−*^) of 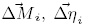, and *ϕ_j_*, and *f* (*ϕ_j_|s_ϕ_*) is the prior for the protein synthesis rate *ϕ_j_* ~ LogN(*m_ϕ_, s_ϕ_*). In order to enforce the restriction that *E*[*ϕ_j_*] = 1 for all genes we include the constraint that 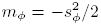. As a result there is only one free parameter for the distribution *f* (*ϕ_j_|s_ϕ_*). Further, we propose a flat prior for s_*ϕ*_, i.e. *f* (*s_ϕ_*) = 1 for s_*ϕ*_ *>* 0.

Figure 9 presents an overview of the structure of our approach, but to summarize,

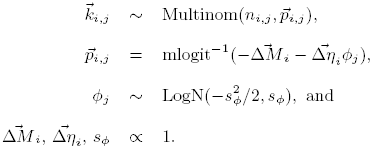

Our MCMC routine provides posterior samples of the genome wide parameters 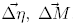, and s_*ϕ*_ and the gene specific, protein synthesis parameters 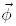. We refer to this model as the ROC SEMPPR *without 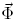* model.

**Figure 9:**
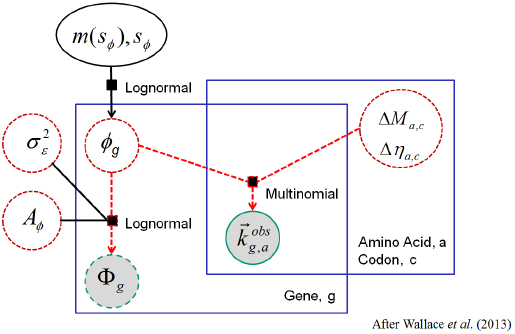
Dependence graph of *with* and *without* ROC SEMPPR methods. Shaded circles 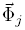 and *k_i,j_* represent observed data. Dashed circles represent key random model parameters while the solid oval represents a random hierarchical parameter. Solid black squares provide information on the distributional relationships between quantities. Large rectangular boxes represent replication of each model component across both amino acids and genes, e.g. pausing, and mutation parameters differ across amino acids but are common across genes, while counts *k_i,j_* differ across both amino acids and genes.

**(Approximate Location of Figure 9)**

We refer to the more general model which incorporates information on *ϕ_j_* from noisy protein synthesis measurements or their proxy, such as mRNA abundances, as the *with 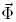* model. This model differs from that of Wallace *et al*. (2013) in that (a) we assume *ϕ_j_* is drawn from a log-normal distribution rather than an asymmetric Laplace distribution, (b) we include and estimate an explicit empirical scaling term *A*_Φ_ for the 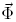 data, and (c) as in the *without 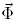* approach, we force the prior for *ϕ_j_*, *f* (*ϕ_j_|s_ϕ_*), to have *E*[*ϕ_j_*] = 1 instead of rescaling estimates of *ϕ_j_* as a post-processing step. This prevents the introduction of additional biases in our parameter estimates. See the Supporting Materials for more details.

#### Model Fitting Details

We briefly describe the model fitting procedure here; full details can be found in Chen *et al*. (Prep). The code was originally based on a script published by Wallace *et al*. (2013), which was modified extensively and expanded greatly. Unless otherwise mentioned, all model fits were carried out using R version 3.0.2 (R Core Team, 2013) using standard routines, specifically developed routines, and custom scripts. All code was run on a multicore workstation with AMD Opteron 6378 processors. For both ROC SEMPPR’s *with* and *without* model fits, it takes *<*30 min and less than 3GB of memory to run 10,000 iterations of a chain when using 5,346 genes of *S. cerevisiae* S288c genome. Each MCMC sampling iteration was divided into three parts:

1. conditional on a new set of parameters, propose new 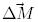 and, 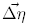 values independently for each amino acid,
2. conditional on the updates of (1), propose a new s_*ϕ*_ value for the prior distribution of 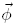, and
3. conditional on the updates of (2), propose new 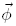 values independently for each gene. Update the new set of parameters and return to (1).

In all three phases, proposals were based on a random walk with step sizes normally or log-normally distributed around the current state of the chain.

In order to generate reasonable starting values for 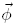 in the *without 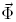* version of ROC SEMPPR, we first calculated the SCUO value for each gene (Wan *et al*., 2006) and then ordered the genes according to these corresponding values. We then simulated a random vector of equal dimension to 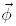 from a 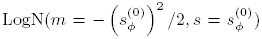 distribution where 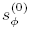 represents the initial value of *s*_*ϕ*_ and controls the standard deviation of *ϕ*. Next, these random 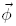 variates were rank ordered and assigned to the corresponding gene of the same SCUO rank. As a result, the rank order of a gene’s initial *ϕ_j_* value, *ϕ_j_,*0, was the same as the rank order of its SCUO value. We tried a variety of 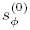 values and they all converged to similar parameter values. For the *with 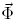* model, we tried both the SCUO based approach and using the 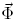 data to initialize our values of *ϕ*. In this second scenario, we set 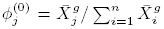 where 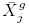represents the geometric mean of the observed mRNA abundances for gene *j*. As in the *without 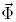* ROC SEMPPR model fit, we found the *with 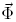* chains consistently converged to the same region of parameter space independent of the initial *ϕ* values. It is worth noting that the structure of the probability function defined in Equation (7) is such that if the rank order of 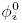 were reversed from their true order, the model would converge to a similar quality of model fit and the signs of the parameters would change. Thus it is recommended that model fits be checked to ensure that the final estimates of *ϕ* for housekeeping genes, such as and ribosomal proteins, are much greater than 1.

Treating our initial protein synthesis rates *ϕ* for the entire genome as explanatory variables, the initial values for 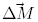 and 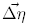 were generated via multinomial logistic regression using the `vglm()` function of the **VGAM** package (Yee, 2013). We also used the covariance matrix returned by `vglm()` as the proposal covariance matrix for 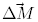 and 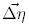for each amino acid. In order to make our random walk more efficient, we used an adaptive proposal function for all parameters in order to reach a target range of acceptance rates between 20 and 35%. For example, the covariance matrix of the step sizes was multiplied by a scalar value that was then increased or decreased by 20% every 100 steps when the acceptance rate of a parameter set was greater than 35% or less than 20%, respectively. The variance terms of the random walks for the 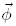 and the global parameter s_*ϕ*_ were also adjusted in a similar manner.

The results presented here were generated by running the MCMC algorithm for 10,000 iterations and, after examining the traces of the samples for evidence of convergence, selecting the last 5,000 iterations as our posterior samples. The arithmetic means of the posterior samples were used as point estimates based on the mean of our posterior samples. Posterior credibility intervals (CI) are generated by excluding the lower and upper 2.5% of samples. Additional details on the model fit can be found in the Supporting Materials and in (Chen *et al*., Prep). The code is implemented in an R package **cubfits** (Chen *et al*., 2014) which is freely available for download at http://cran.r-project.org/package=cubfits.

### Estimating Selection Coefficients using FMutSel

In order to evaluate the consistency of our estimates of *S* = *−*Δ*η ϕ* with other approaches, we used the dataset from Rokas *et al*. (2003) which consisted of 106 aligned genes from 8 yeast species. Details of the model fitting can be found in Kubatko *et al*. (view)(available at bioRxivdoi:http://dx.doi.org/10.1101/007849), but briefly, we used the maximum likelihood tree found by Rokas *et al*. (2003) and then generated MLEs of of the stationary probability of a given codon under the FMutSel model from Yang and Nielsen (2008) using CODONML in PAML 4.4 (Yang, 2007). Using the same notation as in Yang and Nielsen (2008) we have,

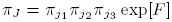

where, for a given gene, *π_J_* represents the stationary probability of observing codon *J* given nucleotide specific mutational bias terms *π*_*j*1_, π_*j*2_, and *π* _*j*3_ and where *F* = ln(Fitness)2*N*_*e*_. It follows that the comparable selection coefficients on synonymous codon usage relative to our reference codon 1 is,

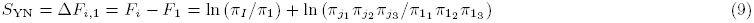

A list of these parameter estimates can be found in the Supporting Materials.

## Acknowledgments

We wish to acknowledge financial support for this project from NSF grants MCB-1120370 (M.A.G. and R.Z.) and EOB (Brian O’Meara, M.A.G., and R.Z.). Additional support was also provided by the National Institute for Mathematical and Biological Synthesis (NSF:DBI-1300426 with additional support from the University of Tennessee). We are grateful to the the RDAV group at the National Institute for Computational Sciences: George Ostrouchov, Drew Schmidt, and Pragnesh Patel who contributed to an earlier attempt to address this problem. We would also like to thank W. Preston Hewgley, Brian O’Meara, Ivan Erill, and Patrick O’Neill for their helpful discussions and suggestions and Laura Kubatko for providing the FMutSel output. Finally, we like to thank our two anonymous reviewers whose comments and suggestions greatly improved the quality of this article.

## Supporting Materials

Supporting Materials for *Estimating gene expression and codon specific translational efficiencies, mutation biases, and selection coefficients from genomic data alone* by Gilchrist *et al*. (In Review).

### Model Validation using Simulated Data

In order to verify the reliability of the *with* and *without* 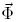 ROC SEMPPR model fits we apply both methods to simulated data. Data set $_1_, is generated from a model with *ϕ* values following a LogN distribution while $_2_ uses the estimates of *ϕ* obtained from our analysis of the S288c genome with 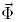 data.

Analysis of both simulated datasets show that both the *with* and *without* 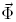 methods produce accurate and unbiased estimates of the mutation bias parameters 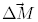 under all circumstances (*ρ* > 0.99, Figures S1 & S2, panels c & d). We also obtained accurate estimates of differences in ribosome pausing times 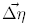. Both *with* and *without* 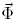 ROC SEMPPR model fits produced near perfect recovery of 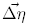 parameters when applied to simulated dataset $_1_ (*ρ* > 0.99, Figure S1, panels a & b).

**Figure S1:**
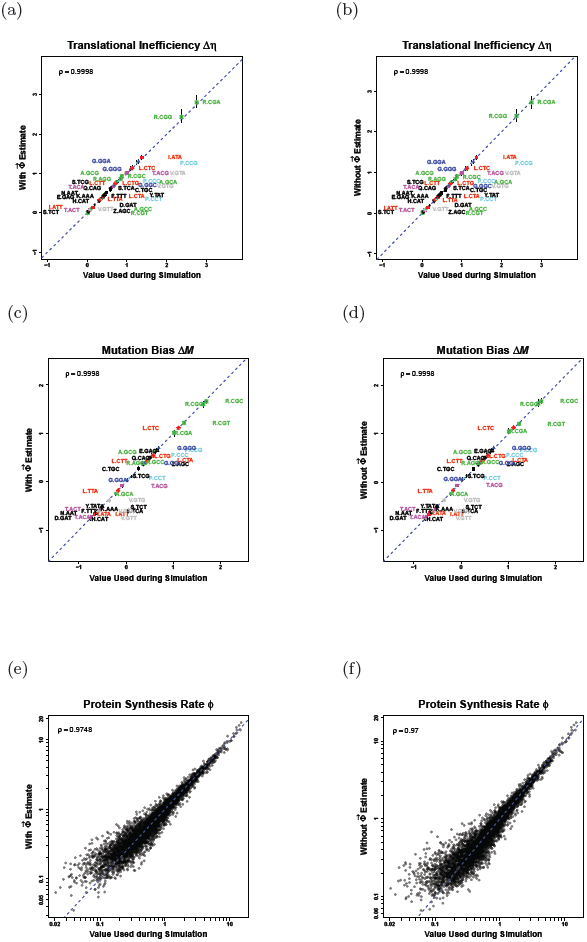
Comparison of estimated parameters versus actual parameters used to simulate data under the model ROC SEMPPR. Here *ϕ ∼* LogN as assumed when fitting ROC SEMPPR. (a) Comparison of *with* 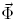 ROC SEMPPR parameter estimates Δ*η* vs. actual data generating parameters Δ*η*. (b) Comparison of *without* 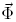 ROC SEMPPR parameter estimates Δ*η* vs. actual data generating parameters Δ*η*. (c) Comparison of *with* 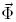 ROC SEMPPR parameter estimates Δ*M* vs. actual data generating parameters Δ*M*. (d) Comparison of *without* 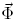 ROC SEMPPR parameter estimates Δ*M* vs. actual data generating parameters Δ*M*. (e) Comparison of *with* 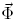 ROC SEMPPR parameter estimates *ϕ* vs. actual data generating parameters *ϕ*. (f) Comparison of *without* 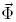 ROC SEMPPR parameter estimates *ϕ* vs. actual data generating parameter *ϕ.*

**Figure S2:**
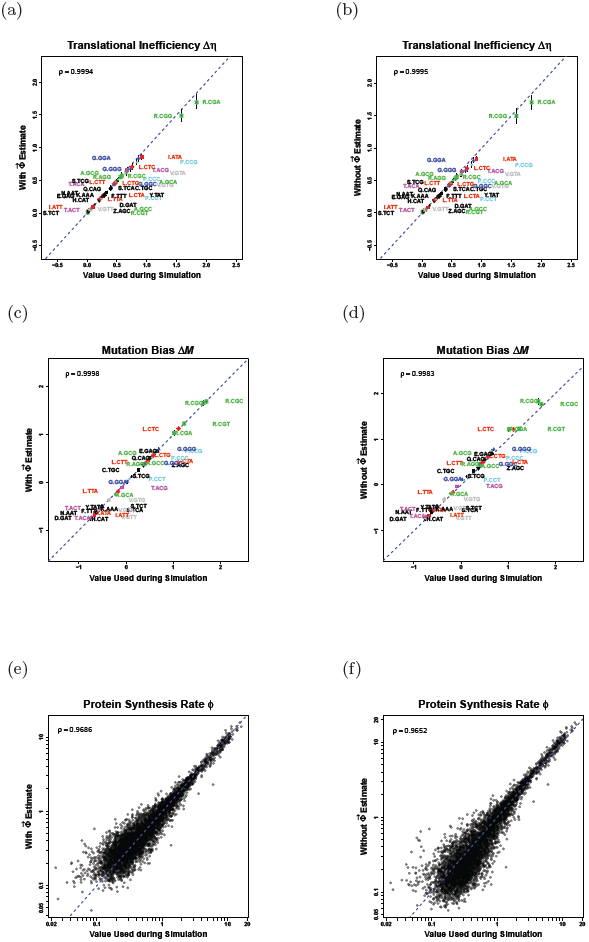
Comparison of estimated parameters versus actual parameters used to simulate data under the model ROC SEMPPR. Here *ϕ* values used in the simulation were based on the *with* 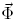 fit of the *S. cerevisiae* S288c genome dataset and, as a result, do not follow a log-normal distribution as assumed when fitting ROC SEMPPR: (a) Comparison of *with* 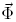 parameter estimates Δ*η* vs. actual data generating parameters Δ*η*. (b) Comparison of *without* 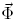 parameter estimates Δ*η* vs. actual data generating parameters Δ*η*. (c) Comparison of *with* 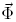 parameter estimates Δ*M* vs. actual data generating parameters Δ*M*. (d) Comparison of *without* 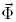 parameter estimates Δ*M* vs. actual data generating parameters Δ*M*. (e) Comparison of *with* 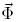 parameter estimates *ϕ* vs. actual data generating parameters *ϕ*. (f) Comparison of *without* 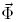 parameter estimates *ϕ* vs. actual data generating parameters 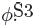

**Figure S3:**
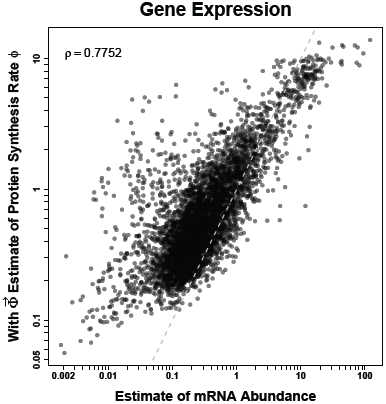
Comparison between posterior mean estimates of *ϕ* for the *with* 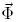 model fit and 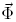 data consisting of mRNA abundance measurements from Yassour *et al*. (2009).

When applied to simulated dataset $_2_, both *with* and *without* 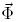 estimates of 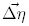 showed strong agreement with parameter values (*ρ* > 0.99, Figure S2, panels a & b). We did, however, observe a small downward bias in their absolute values (∼ 7%). This is a special case of attenuation bias (Fuller, 1987) which results from the *ϕ* values in $_2_ being distributed with a heavier right tail than the corresponding LogN distribution with the same mean and variance.

Comparing the *with* and *without* 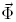 ROC SEMPPR estimates of protein synthesis rates, e.g. the posterior means, 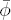 and the *ϕ* values used in our simulations illustrates the predictive power of ROC SEMPPR. For example, analysis of the simulated dataset $_1_ indicates that under ideal conditions we observe correlation coefficients between the log of our protein synthesis estimates, log(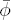), and the log of their true values, log(*ϕ*) of *∼* 0.96 for both the *with* and *without* 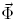 ROC SEMPPR model fits (Figure S1). Even when the true distribution of *ϕ* values violates the LogN assumption as in $_2_, we still observe correlation coefficients between log(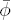) and log(*ϕ*) of *∼* 0.96 (Figure S2).

### Scaling Bias due to Noise and Inherent Uncertainty

Because measurements of mRNA abundances, whether via microarray florescence or sequencing data, are usually not scaled to any particular unit, researchers often use either the sum of all the measurements or their mean value as a means of scaling their results. While it is intuitive to scale the data in this way, if the additional measurement noise is not taken into account a subtle biases on *ϕ* and 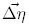 is introduced. The nature of the bias can be most easily illustrated when we assume that both the signal and the noise follow log-normal distributions, however, the effects should be present as long as the noise is not symmetrically distributed around the underlying signal values.

For example, let 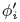 represent the true, unscaled protein synthesis rate of gene *i*, i.e 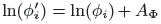 and assume that, across the genome, 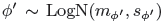, such that 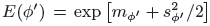 Let ϕ_*i,j*_ represent a given noisy observation or estimate of 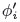, i.e. ϕ_*i,j*_ is part of our 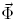 data set. Also let 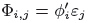 where *ε_j_ ∼* LogN(0, *s*_*ε*_) and implies that the observation ϕ_*i,j*_ is log normally distributed around the true values 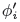. Even though the noise is centered around the true value, because the log-normal distribution is asymmetric, 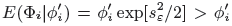 and when considering the entire distribution 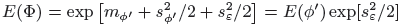. Thus we see that the mean of our observed values is actually greater than the mean of the true signals underlying them and, as a result, if one scales by the sum or the mean of these observed values the resulting values will be biased downward by a factor of 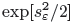. To remove this bias, we introduce an additional scaling term *A*ϕ such that 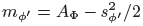 and, as a result, *E*(*ϕ′*) = exp [*A*_ϕ_] and 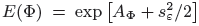. Our empirical data provides an estimate of *E*(ϕ) and the inconsistency between the degrees of adaptation in CUB observed across genes and their expression levels greater than that expected due to genetic drift allows us to estimate *s*_*ε*_ while, simultaneously estimating *A*_ϕ_.

Finally, we note that simply scaling one’s estimates of *x* by the mean of these estimates during the MCMC run also introduces bias. This is because our estimates of 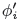 during the MCMC, ϕ_MCMC_ are imprecise and, as a result, their mean value will be overestimated. Assuming our uncertainty in *x* is lognormally distributed LogN(*m* = 0, *s* = *s*_MCMC_), 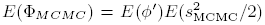. As a consequence, the scaled protein synthesis rates, *ϕ*, are biased downward leading to an overestimation in the absolute differences in pausing times between codons, 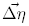 The effects of this bias are actually evident in Wallace *et al*. (2013) Figure 5A where the estimates of the coefficients differ from the values used during their simulations. Including the parameter *A*_ϕ_, which explicitly models this scaling terms, provides a simple way to avoid these issues.

### Fitting of Model to Genomic Data and Noisy Measurements of Protein Synthesis

We generalize our ROC SEMPPR model to include the extraction of information from noisy, unscaled measurements of protein synthesis for each gene, i.e. 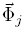. This is essentially the same model as Wallace *et al*. (2013) except instead of rescaling estimates of 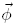 and 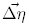 in preand post-MCMC data processing step, we include the estimation of the scaling term *A*_ϕ_ discussed in the last section.

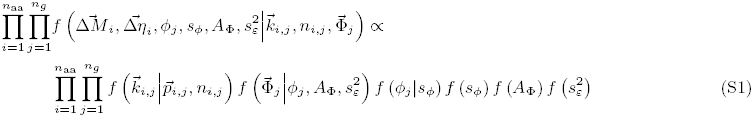

where, as before, 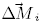 and 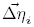 are the mutation and selection coefficients respectively for amino acid *i*, 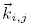 are the codon counts following a multinomial distribution for the amino acid *i* in the ORF of gene *j* as defined in Equation (2), *n*_*i,j*_ is the sum of all codon counts related to a particular amino acid *i* in the gene *j*, 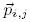 is an inverse multinomial logit function of 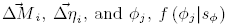, is the prior for the protein synthesis rate 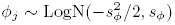, and *f*(*s*_*ϕ*_) = 1.

Additionally, we assume that log 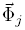 ~ N (log (*ϕ_j_*) + *A*_ϕ_, 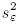, i.e. the log transformed measurements log 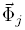 are offset by a constant *A*_ϕ_ and normally distributed around log(*ϕ*) + *A*_ϕ_ with variance 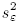. We also assume *f*(*A*_Φ_) = 1 and 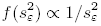. Both *A*_ϕ_ and 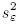 are genome scale parameters and are estimated in the *with* 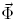 model. In the future, the assumption that 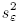 is the same across genes could be relaxed. In the absence of any 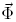 data, the *f*(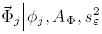), *f* (*A*_Φ_), and *f*(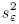) terms are undefined and drop out.

The system below summarizes the expressions just given describing Equation (S1):

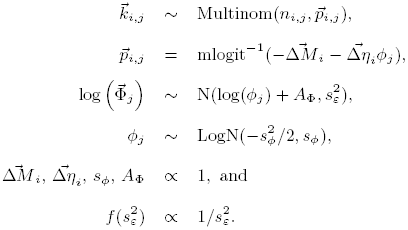

To fit the *without* and *with* 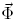 models, we apply the following algorithm with a superscript (*i*) indicating the *i*^th^ iteration of an MCMC chain.

Step 1. Update 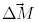 and 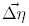 conditional on all other parameters in the *i*^th^ iteration through a random walk Metropolis-Hasting (MH) algorithm:

a. Step *i* = 0 only.

i. Calculate SCUO value for each gene following Wan *et al*. (2006).
ii. Generate random ordered values *ϕ*^(0)^ by simulating from 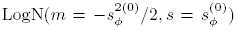, and sorting them in the same order as the SCUO values to maintain the rank order of production rates among genes.
iii. Given *ϕ*(^0^), for each amino acid *a* estimate initial values 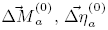, and the covariance matrix of these estimates 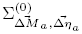 using multinomial logistic regression.
b. For each amino acid, independently simulate a new proposal for 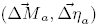 jointly from a multivariate normal distribution which has mean 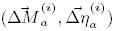) and covariance 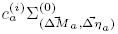 with initial adaptive scaling factor 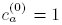. See Marin and Robert (2007, Chapter 2) for details on incorporating a covariance matrix in practice.
c. Accept the proposal with the MH probability based on the acceptance ratio and set 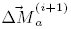 and 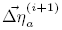 accordingly for all amino acids.
Step 2. Update hyperparameters conditional on all other parameters:
  a. If using the fitting *with* 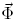 model: update 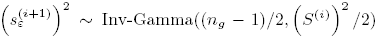 where 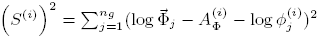.
  b. Update 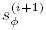 using a random walk MH with proposal distribution 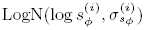 with initial value 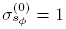 for the adaptive scaling factor of MCMC. Also, set 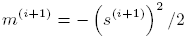.
  c. If fitting *with* 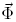 model: update 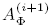 using a random walk MH with proposal distribution 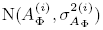 with initial value 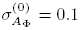 for the adaptive MCMC scaling factor.

Step 3. Update protein translation rates conditional on Steps 1 and 2 and all other parameters: For each gene *j*, generate *ϕ_j_* through a random walk MH step:

a. Propose *ϕ_j_* from 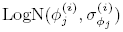 with initial value 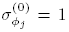 for the adaptive MCMC scaling factor.
b. Accept the proposal with the MH probability based on the acceptance ratio and set 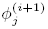 accordingly.

Step 4. Update all adaptive scaling factors if the acceptance rate of each set of parameters falls outside the 20-35% acceptance rate in the above Steps 1, 2, and 3 in order to sample the posterior distribution efficiently.

### Comparison of Predicted Protein Synthesis Rates *ϕ* to Independent mRNA Abundance Measurements

Figure S4 compares posterior mean estimates of *ϕ* produced *with* (using the mRNA abundance measurements of Yassour *et al*. (2009)) and *without* 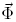 to four additional lab measurements of mRNA abundances reported by Arava (2003); Nagalakshmi *et al*. (2008); Holstege *et al*. (1998); Sun *et al*. (2012). These values can be found in Table S9. Correlation coefficients are provided for each figure and tend to be slightly higher for estimates generated using the *with* 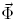 algorithm. Although this seems to indicate that *with* 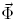 estimates are superior, it is worth noting that these data measure mRNA expression levels. Because the *without* 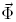 algorithm estimates protein synthesis rates, fundamentally a different quantity, we would expect these estimates to differ. Because the *with* 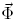 measurement algorithm shrinks the protein synthesis estimates toward the mRNA expression observations, it is natural that *with* 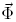 estimates show higher correlation with measurements from other laboratories.

**Figure S4:**
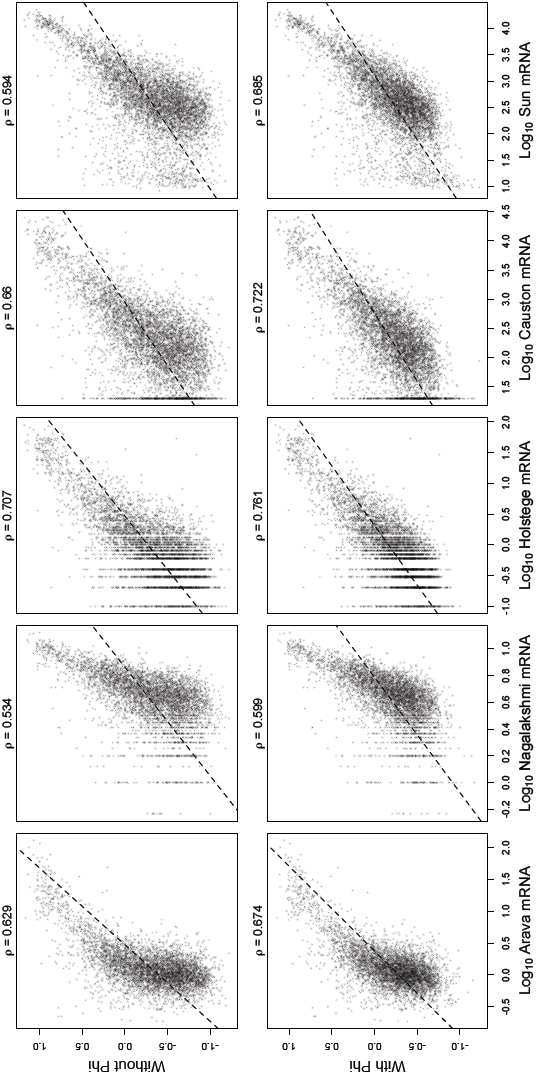
Scatter plot comparisons of *with* (Yassour measurements) and *without* 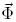 posterior mean estimates to empirical measurements from four additional laboratories. The units for *ϕ* are protein/*t* and time is scaled such that the prior for *ϕ* satisfies *E*(*ϕ*) = 1. The empirical mRNA abundance measurements, [mRNA], are being used here as a proxy for protein synthesis rates, i.e. [mRNA] ∝ protein/*t*. The measurements are scaled such that the mean [mRNA] value is 1. Pearson correlation coefficients *ρ* are given and the dashed black line represents thet of a linear regression model.

**Figure S5:**
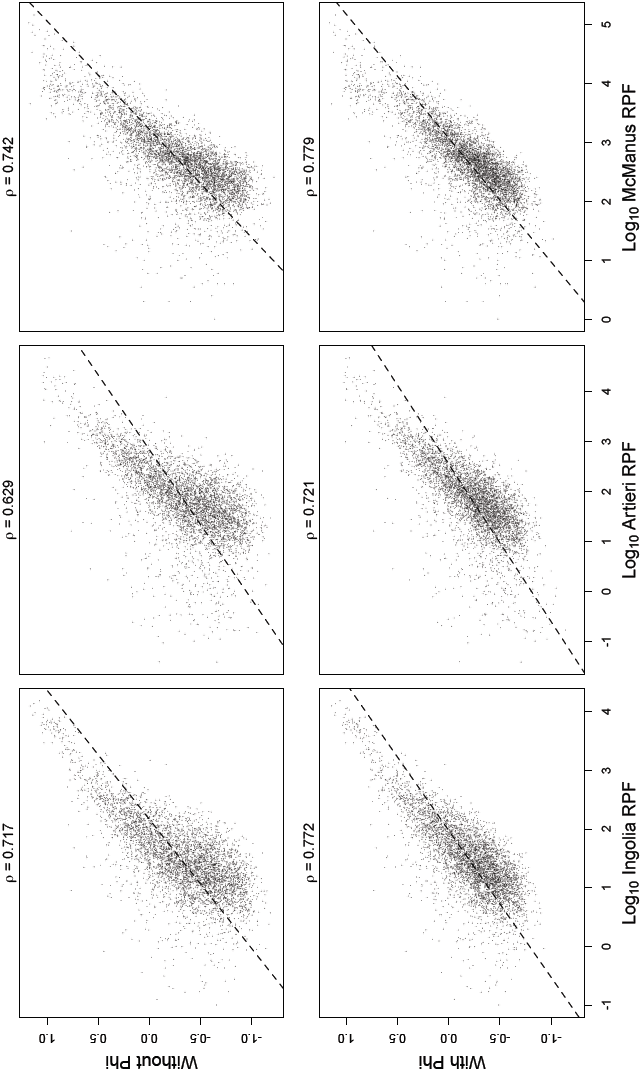
Scatter plot comparisons of *with* 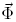 (Yassour) and *without* 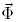 posterior mean estimates to empirical measurements from three ribosome profiling datasets from three different laboratories. The units for *ϕ* are protein/*t* and time is scaled such that the prior for *ϕ* satisfies *E*(*ϕ*) = 1. The empirical ribosome profiling measurements were originally in units of reads per kilobase of transcript per million mapped (rpkm) corrected for mRNA length. These measurements are scaled such that the mean rpkm value is 1. Pearson correlation coefficients ρ are given and the dashed black line represents thet of a linear regression model.

### Supplemental Tables

Data in supplemental tables can be downloaded from **doi:** http://dx.doi.org/10.1101/009670

S1. Summary statistics of posterior estimates of Δ*M* for *S. cerevisiae* S288c genome estimated *with* 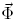 (s288c deltam wphi.tsv).

S2. Summary statistics of posterior estimates of Δ*M* for *S. cerevisiae* S288c genome estimated *without* 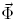(s288c deltam wophi.tsv).

S3. Summary statistics of posterior estimates of Δ*η* for *S. cerevisiae* S288c genome estimated *with* 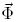 (s288c deltaeta wphi.tsv).

S4. Summary statistics of posterior estimates of Δ*η* for *S. cerevisiae* S288c genome estimated *without* 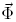 (s288c deltaeta wophi.tsv).

S5. Summary statistics of posterior estimates of *ϕ* for *S. cerevisiae* S288c genome estimated *with* 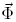(s288c phi wphi.tsv).

S6. Summary statistics of posterior estimates of *ϕ* for *S. cerevisiae* S288c genome estimated *without* 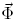 (s288c phi wophi.tsv).

S7. Gene and codon specific selection coefficients for *S. cerevisiae* S288c genome estimated *with* 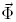 (s288c selection coefficient wphi.tsv).

S8. Gene and codon specific selection coefficients for *S. cerevisiae* S288c genome estimated *without* 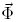 (s288c selection coefficient wophi.tsv).

S9. Additional absolute mRNA measurements from multiple laboratories of *S. cerevisiae* Genome (s.cerevisiae.mRNA.measurements.tsv).

S10. Additional measurements of protein synthesis rates from ribosome profiling experiments from multiple laboratories of *S. cerevisiae* Genome (s.cerevisiae.rpf.measurements.tsv).

S11. Results from linear regression of FMutSel estimates of *S* vs. *without* 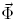 ROC SEMPPR estimates of *S* for the 106 genes in the Rokas *et al*. (2003) dataset (FMutSel S vs ROC wo phi S regressions.txt).

